# Modular RNA motifs for orthogonal phase separated compartments

**DOI:** 10.1101/2023.10.06.561123

**Authors:** Jaimie Marie Stewart, Shiyi Li, Anli Tang, Melissa Ann Klocke, Martin Vincent Gobry, Giacomo Fabrini, Lorenzo Di Michele, Paul W.K. Rothemund, Elisa Franco

**Affiliations:** Department of Computing and Mathematical Sciences, California Institute of Technology; Pasadena, 91125, CA, USA; Department of Bioengineering, University of California at Los Angeles; Los Angeles, CA 90024, USA; Department of Mechanical and Aerospace Engineering, University of California at Los Angeles; Los Angeles, CA 90024, USA; Interdisciplinary Nanoscience Center (iNANO), Aarhus University; Aarhus, DK-8000, Denmark; Department of Chemistry, Molecular Sciences Research Hub, Imperial College London; London W12 0BZ, UK; Department of Chemical Engineering and Biotechnology, University of Cambridge; Cambridge CB3 0AS, UK; fabriCELL, Molecular Sciences Research Hub, Imperial College London; London W12 0BZ, UK

## Abstract

Recent discoveries in biology have highlighted the importance of protein and RNA-based condensates as an alternative to classical membrane-bound organelles for the task of compartmentalizing molecules and biochemical reactions. Here, we demonstrate the rational design of pure RNA condensates from star-shaped RNA motifs. We generate condensates using two different RNA nanostar architectures: multi-stranded nanostars whose binding interactions are programmed via single-stranded overhangs, and single-stranded nanostars whose interactions are programmed via kissing loops. Through rational design of the nanostar interaction sequences, we demonstrate that both architectures can produce orthogonal (distinct and immiscible) condensates, which can be individually tracked via fluorogenic aptamers. We also show that aptamers make it possible to recruit peptides and proteins to the condensates with high specificity. Successful cotranscriptional formation of condensates from single-stranded nanostars suggests that they may be genetically encoded and produced in living cells. We provide a library of orthogonal RNA condensates that can be modularly customized and offer a route toward creating systems of functional artificial organelles.

---

The discovery of membraneless organelles is transforming our understanding of cellular biology and disease^1^. These organelles, also known as biomolecular condensates, arise when mixtures of nucleic acids and proteins segregate into spatially separated phases due to specific and non-specific (electrostatic and hydrophobic) molecular interactions^2^. Condensation broadly describes the formation of viscoelastic aggregates, and in biology it is typically associated with a phase transition of mixtures of both RNA and proteins^2,3^ that is driven by the interaction of intrinsically disordered domains (IDRs) of proteins^4^, protein-RNA interactions^5^ or RNA-RNA interactions^6^. Distinct, immiscible condensates — here termed “orthogonal condensates” — arise through variations of the chemical identity and interactions of the protein and RNA components from which the condensates are composed. The prevailing model is that orthogonal condensates enable the spatial and temporal control of chemical components and biochemical reactions. In general, the greater the number of orthogonal condensates that can be created, the greater the number of distinct biochemical functions that can be performed. In cells there are dozens of functionally distinct condensates^7^, that are involved in diverse gene regulation processes^8^, cellular stress^9^, and neurodegenerative diseases such as Alzheimer’s Disease and ALS^10^. The development of libraries of synthetic, orthogonal condensates would allow for the engineering of coordinated systems of controllable microcompartments with distinct functions matching the complexity of living systems.

As sequence-programmable biopolymers, proteins and RNA have all shown promise as building blocks of artificial condensates. A fundamental design principle used in these investigations has been to introduce weak, non-specific, homotypic interactions that are thought to be essential for phase separation. This was made possible by engineering proteins to include IDRs present in naturally condensing proteins like FUS and SUMO proteins^11–14^. Similarly, artificial RNA condensates have been demonstrated using long molecules featuring expanded repeats of short sequence^15,16^, which are found in nuclear foci associated with neurological diseases. While IDRs and short repeats introduce multivalency, they also introduce promiscuous molecular interactions, thereby limiting the possibility of building coexisting yet immiscible condensates with distinct identities and tunable properties. An alternative approach has been pioneered through nanostructured DNA motifs that form condensates thanks to localized interactions whose specificity is achieved by rationally designed base-pairing^17–19^. DNA nanostars have successfully been used to build coexisting but immiscible orthogonal condensates with programmable phase transitions^17,19,20^.

Taking inspiration from the success of DNA nanostar condensates, and using techniques from RNA nanotechnology^21,22^ we have designed and synthesized both multi-stranded and single-stranded RNA motifs that phase separate into orthogonal condensates. With the aid of computational models predicting RNA-RNA interactions, we demonstrate a suite of star-shaped RNA motifs, or nanostars, that generate RNA-dense droplets thanks to designed base-pairing domains, either linear sticky-ends or kissing loops, at the tip of their arms (Fig. 1). As shown in DNA nanostar analogues, this strategy makes it possible to obtain phase transitions by controlling nanostar affinity (determined by sequence and length of the base-pairing domains) and valency (determined by the number of arms)^17,18,23^. We adapt RNA nanostars to include an array of RNA domains that bind to small molecules and peptides, producing an expandable library of modular motifs that produce condensates with the capacity to recruit and segregate client molecules. Finally, we show that single-stranded RNA nanostars fold and produce condensates cotranscriptionally^24^, and could serve as genetically encoded building blocks to make RNA organelles with desired biophysical features and with the capacity to concentrate molecular targets. This feature is immediately applicable to building organelles in synthetic cells, as demonstrated in Fabrini et al^25^.

**Figure 1.**
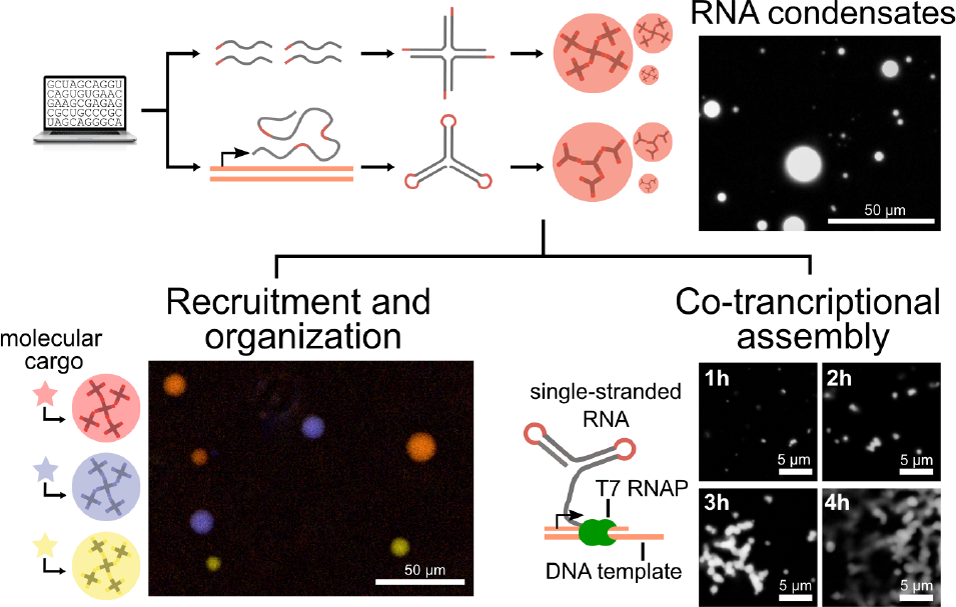
We demonstrate the design and formation of RNA condensates from rationally designed multi-stranded and single-stranded RNA “nanostars.” These condensates have the capacity to recruit and organize molecules, and single-stranded motifs yield condensates cotranscriptionally. Scale bars: 50 μm.

## Results

### Condensates emerge from rationally designed multi-stranded RNA motifs interacting via sticky-ends

We begin by demonstrating RNA nanostars assembled from multiple RNA strands, taking inspiration from DNA nanostars that interact via linear sticky-ends^18^ (Fig. 2A). In our designs, arm sequences are distinct, but each arm presents the same palindromic sticky-end that controls nanostar affinity. We built several four-arm variants, labeled 4m1, 4m2, etc., where “m” denotes the multi-stranded nature of the constructs, “4” is the number of arms, and the final number denotes the sticky-end variant. The sequences of 15-nucleotide (nt) arms and their sticky-ends were optimized using NUPACK^26^ to prevent repeats and unwanted interactions. To improve flexibility, we included unpaired adenines at the nanostar core and at the base of the sticky-ends^27^. RNA strands were suspended in an assembly buffer (40 mM HEPES, 100 mM KCl, 500 mM NaCl), thermally treated by a “melt and hold” protocol consisting of a quick 70°C melt followed by a 12h-long hold at 50°C (Fig. 2B), and stained with 1X SYBR gold for microscopy. We will refer to this buffer and thermal treatment as our standard experimental conditions. Additional hold temperatures and buffer conditions were screened in the Supplementary Information file (Fig. S1-S3).

**Figure 2:**
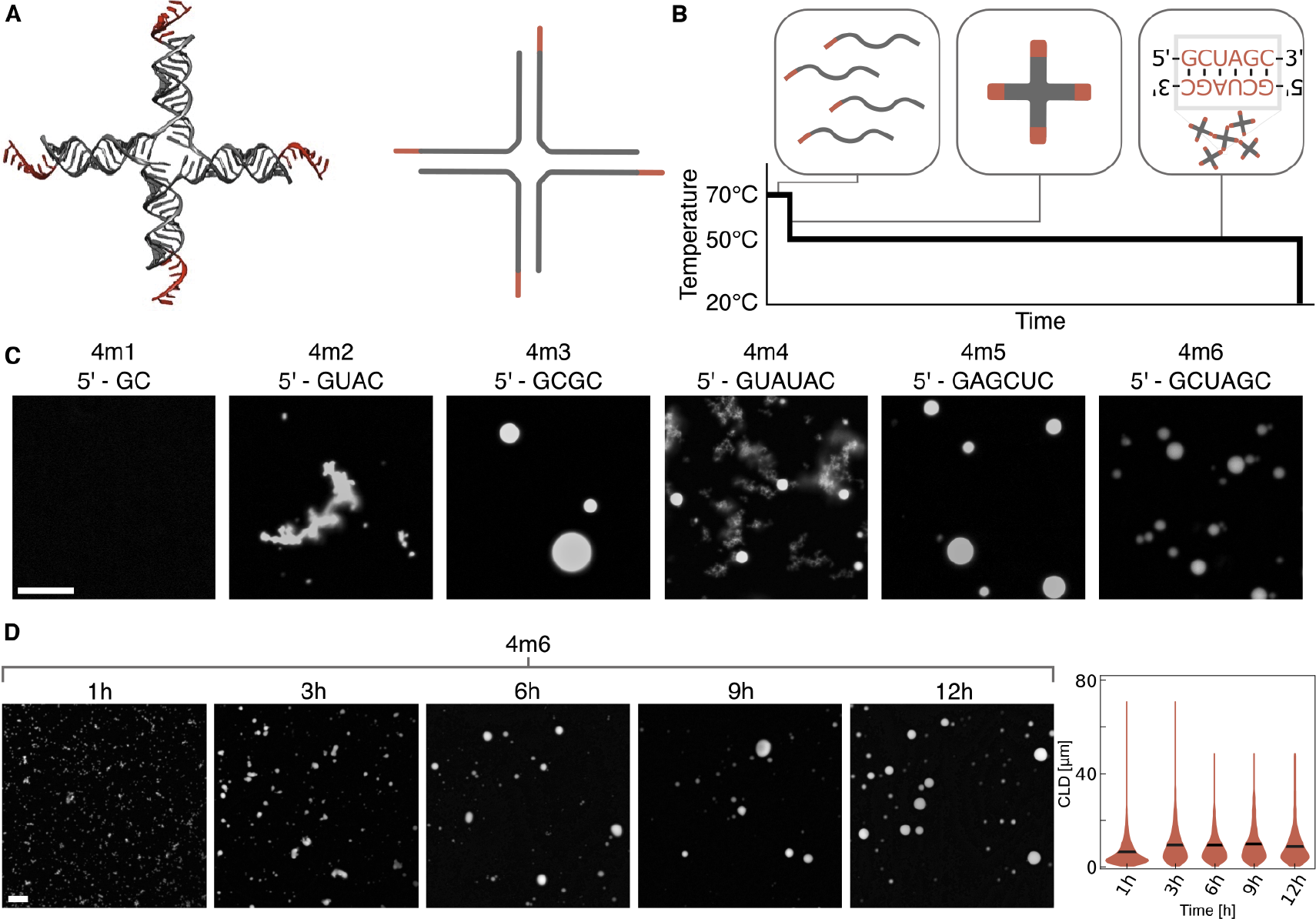
Multi-stranded RNA nanostars yield condensates of variable size and morphology. (A) PDB rendering and 2D representation of a multi-stranded RNA nanostar. (B) We use a “melt and hold” thermal annealing that sequentially promotes RNA denaturation (left), nanostar assembly (middle), and nanostar-nanostar interactions (right) that yield condensates. (C) Condensate morphology depends on sequence and length of nanostar sticky-ends. (D) Growth of multi-stranded condensates (4m6) formed in 40 mM HEPES/100 mM KCl/500 mM NaCl and sampled for imaging over a 12-hour incubation period. Right: condensate growth during incubation is confirmed by the image chord length distributions (CLD); black lines indicate the mean. Samples were stained with SYBR Gold and imaged. Images are representative of data collected in three replicates. Violin plots in (D) pool data from one sample, imaged 14 times. Scale bars: 40 μm.

We found that 4-6 nt long sticky-ends yield condensates of varying morphology, including round droplets, aggregates of slowly fusing droplets, and “cloudy” aggregates (Fig. 2C). In contrast, no condensates formed in the absence of sticky-ends or with 2 nt long sticky-ends unless excess salt was added^27^ (Fig. 2C and Fig. S4). We observed the presence of gel-like aggregates in sticky-ends with high UA content. We tracked condensation of variant 4m6 during the temperature hold, finding large condensates after 3-6 hours of incubation (Fig. 2D). The average size of condensates increases during the hold step, as shown via chord length distribution (CLD) analysis^28–31^ (Fig. 2D, right, and SI Methods 1.4), and their spherical shape is indicative of a liquid-like state. We tested the influence of the hold temperature on variant 4m5, finding that lower hold temperatures (40-45°C) yield aggregates, while temperatures in the range of 46-65°C produce spherical liquid-like assemblies; a 70°C hold eliminates condensation (Fig. S1). Condensates persist at high KCl concentrations (Fig. S2), and the addition of MgCl_2_ alters their melting temperature (Fig. S3). We finally verified that a three-arm nanostar variant (adapted from variant 4m6) also yields condensates in our standard conditions (Fig. S5), consistent with previous work showing that three arms are sufficient to promote condensation of DNA nanostars^17^.

### Loop-loop interactions enable condensation of single-stranded RNA motifs

A requirement for potentially producing RNA nanostars in living cells is that they form at constant temperature, as they are being transcribed. To achieve this, we developed RNA nanostars comprising a single strand rather than several distinct molecules. In these designs, sticky-ends are replaced by kissing loop (KL) domains placed at the end of consecutive stems that serve as nanostar arms (Fig. 3A). Like in the multi-stranded designs, the arm sequences are distinct to minimize misfolding but all KLs on a particular nanostar are identical. Given our goal of cotranscriptional formation, we decided to minimize transcript length and include only three consecutive arms, the lowest valency for condensation. Stem sequences were adapted from design 4m6, eliminating one of the arms. We started by testing the palindromic wild-type HIV KL sequence (GCGCGC, variant termed 3sWT, where “s” denotes the single-stranded nature of the design and “3” the number of arms), and we also developed variants with non-palindromic sticky-ends (3sα-ζ), each including 2 distinct nanostars (Fig. 3B). Before attempting cotranscriptional folding, we characterized the condensation of our designs using purified RNA treated with our melt and hold annealing protocol, using assembly conditions consistent with those used for the multi-stranded constructs.

**Figure 3:**
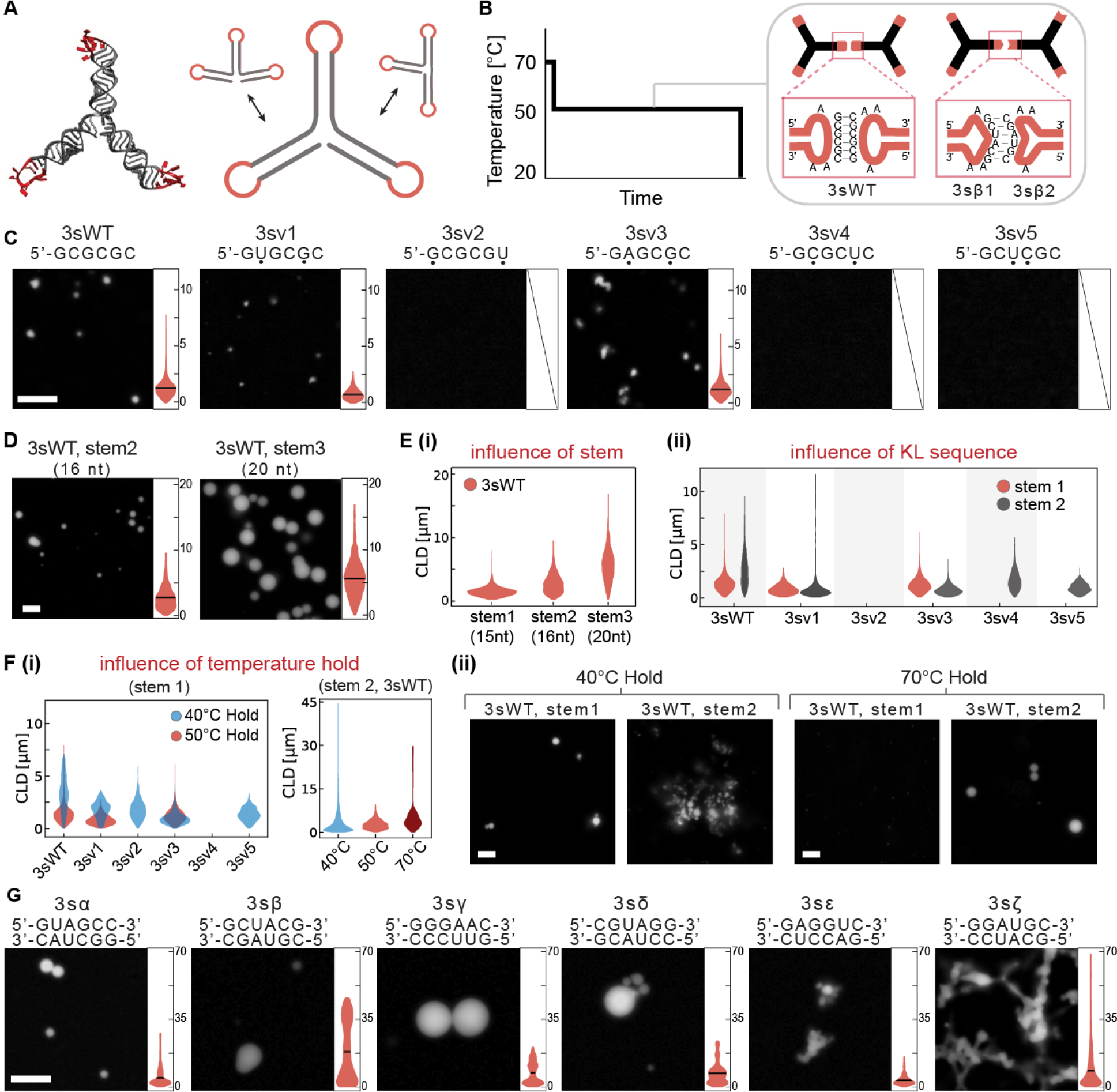
Condensates generated by single-stranded RNA nanostars. (A) PDB and 2D representation of a single-stranded RNA nanostar. (B) Melt and hold thermal annealing protocol; single-nanostar systems with palindromic KL (left) and two-nanostar systems with non-palindromic KL (right) condense during the hold phase. (C) Example images and violin plots of the chord length distribution (CLD) of condensates formed by nanostars differing by KL. Black dots (**·**) indicate mismatches or wobble pairs (D) Example images and CLD of condensates forming under different nanostar stems. (E) Influence of the stem sequence and length: (i) longer stems yield larger condensates; (ii) changing stem sequence impacts condensate formation and morphology across KL variants (F) (i) The hold temperature influences formation of stem1 variants; (ii) example images comparing 3sWT-stem1 and 3sWT-stem2 under a 40° and 70°C hold. (G) Two-nanostar systems produce condensates of varying morphology, as captured by violin plots of the CLD. All experiments were replicated at least three times; images are representative examples. Violin plots pool data from three independent replicates. Scale bars: 20 μm.

We found that condensate formation occurs robustly across designs, although it is influenced by KL and stem sequences. The 3sWT nanostars yield abundant condensates that continue to grow throughout the hold step of the annealing protocol (Fig. S8). We verified that interacting KLs are essential for condensation: replacing only one of the three WT KLs with a poly-A sequence suppresses droplet formation (Fig. S9). We then tested 5 variants of the WT KL (dubbed 3sv1-v5) with mismatches or wobble pairs that modulate the probability of KL dimerization^32^ and therefore change the nanostar affinity (Fig. 3C and Figs. S10). Condensates are observed with the WT, v1, and v3 variants, yielding similar chord length distributions, but not with variants v2, v4, and v5, consistently with previous dimerization tests^32^. While KL v2 is expected to dimerize, it did not yield condensates under these assembly conditions, likely due to the high hold temperature and the lack of divalent cations in our assembly buffer. Next, we examined the influence of nanostar stems on condensate formation. We built nanostars with a 16-nt “stem 2” variant whose sequence is adapted from a well-known design for three-arm DNA nanostars^17^, and a 20-nt “stem 3” variant designed using NUPACK. We obtain condensates with both new stem variants with the WT KL (Fig. 3D), and longer arms correlate with larger condensate size, an effect also observed with DNA nanostars^33^ (Fig. 3E (i)). When combining stem 2 with different KL variants, we find that variants v4 and v5 yield condensates (Fig. 3E (ii)) pointing to the fact that the stem-mediated interactions may play a role in condensation under the melt and hold protocol. This role cannot be simply explained thermodynamically, given the similar free energy (ΔG) of stem 1 and stem 2 formation (NUPACK estimates ΔG =-67.45 kcal/mol for stem 1 and ΔG =-68.53 kcal/mol for stem 2 at 50°C, with comparable GC content flanking the KL).

The choice of hold temperature significantly influences condensation (Fig. 3F). Across KL variants, a 40°C hold facilitates condensation for stem 1 when compared to a 50°C hold. This is evidenced by droplets appearing at 40°C while remaining dispersed at 50°C (variants 3sv2 and 3sv5) or by an increase in droplet size (variants 3sWT and 3sv1), as shown in Fig. 3F(i) (example microscopy images are in Fig. S10). All stem 2 variants, except 3sv2, yield condensates under a 50°C hold (Fig. 3E(ii) and Fig. S10). Strikingly, variant 3sWT-stem2 forms condensates even with a 70°C hold, likely due to interactions enabled by partial stem melting (Fig. 3F(i)). The round shape of these condensates suggests a liquid state, at least during the hold phase (Fig. 3F(ii)).

The addition of MgCl_2_ to the assembly buffer alters the observed phase transitions. In this case, also 3sWT-stem1 nanostars yield condensates under a 70°C hold, and variant 3sv2-stem1 also produces condensates under a 50°C hold (Fig. S11). Other variants produce more non-spherical condensates when compared with the standard assembly buffer including monovalent cations (Fig. S11). This behavior is consistent with the fact that divalent cations like MgCl_2_ generally stabilize nucleic acid assemblies^34^, and may increase the affinity of KL nanostars that otherwise do not condense^35^. At the same time, MgCl_2_ can promote aggregation and kinetic trapping^36^, and in this case, it can compromise the mobility of nanostars in the dense phase resulting in the formation of gelatinous or solid assemblies. This is an important consideration because MgCl_2_ is typically required for *in vitro* transcription, and is a ubiquitous component in protocols for DNA or RNA self-assembly.

Finally, we tested 6 more RNA nanostar variants (3sα-ζ) each comprising 2 distinct nanostars (e.g., 3sα1, 3sα2) with non-palindromic KL designed to be complementary (Fig. 3G). By adopting non-palindromic sequences, we can expand by 64-fold the theoretical sequence design space of a 6 nucleotide KL domain. With the melt and hold protocol and our assembly buffer, we found that all two-nanostar variants generated condensates with variable size and morphology (Fig. 3G). Individual nanostars produced no condensate, except for variant 3sε1 because of partial self-complementarity of the KL domains (Fig. S12). Collectively, our data suggest that KL with more than 4 base pairs have a high likelihood of yielding condensation. If 4 base pairs form, the outcome depends on the overall strength of the interactions (Fig. S13).

### Orthogonal RNA condensates can be programmed to recruit guest molecules

We next demonstrate the potential of RNA nanostars to capture client molecules and recruit them to the RNA dense phase. We focus on condensates produced through the melt and hold protocol using purified RNA.

We began by appending client recruitment domains (aptamers) at the tip of one of the arms of variants 4m3, 4m5, and 4m6 (upstream of the 5’ end of the sticky-end). We first validated this idea through fluorescent light-up aptamers (FLAPs) known as Red Broccoli (appended to variant 4m6 and to a three-arm variant 3m1, shown in Fig. S5), Corn (4m5), and Orange Broccoli (4m3), which all bind to fluorophore DFHO but each results in a distinct emission spectrum (Fig. 4A, Fig. S6). (For ease of visualization, we depict FLAPs using the secondary structure predicted by NUPACK^26^, which does not capture their complex 3D tertiary structure^37^.) Because these variants were designed to maximize orthogonality, we expected each nanostar to form distinct condensates that do not mix with the others. Through microscopy we first verified that these nanostars yield condensates when assembled in isolation in the presence of DFHO (Fig. S6), and then that they do not mix when assembled simultaneously. To quantitatively assess the degree of condensate mixing we plotted a histogram of the arctangent of the pixel intensity ratio FITC/Cy3 (Fig. 4A, bottom right, bottom right, and Fig. S14). The histogram peaks of control samples (individually assembled nanostars) are consistent with those of samples including mixed nanostars, indicating that the aptamers are not colocalized. These histograms highlight that although our nanostars are sequence-designed to remain de-mixed, the excitation and emission spectra of the FLAPs have some overlap (Orange Broccoli 513/562 nm, Red Broccoli 518/582 nm, and Corn 505/545 nm)^37^. Further, Orange Broccoli and Red Broccoli have a high degree of sequence similarity, where mutation at nucleotide position 71 is hypothesized to be the key cause of the difference in fluorescence emission^37,38^. Finally, differences in fluorescence intensity are likely due to the difference in the K_D_ of DFHO (Orange Broccoli ∼230 nM, Red Broccoli ∼206 nM, and Corn ∼70 nM^37,38^).

**Figure 4:**
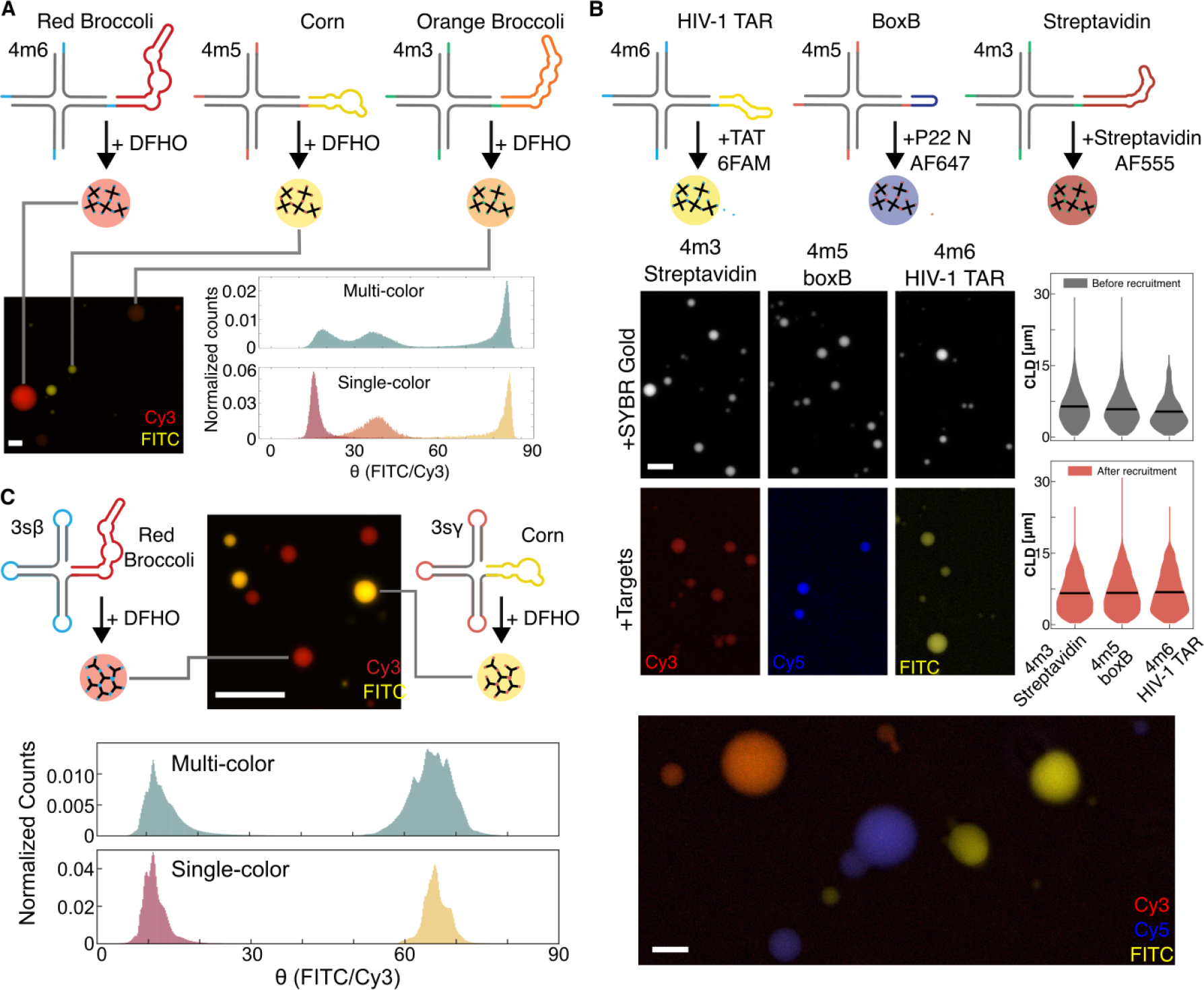
Sequence-orthogonal RNA nanostars produce distinct condensates that recruit clients specifically and remain demixed. (A) Multi-stranded sequence-orthogonal nanostars modified to include FLAPs. When annealed in one pot they produce distinct RNA condensates that do not mix. Condensates were imaged in both FITC and Cy3 channels; each pixel’s FITC/Cy3 ratio was mapped to an angle θ and histogrammed. Histogram peaks for individually-annealed nanostars (single-color) align with those of simultaneously-annealed nanostars (multi-color). (B) Top: Nanostars including aptamer domains that recruit streptavidin, P22 N peptide, and TAT peptide guests. Middle: Example images and violin plots of the chord length distribution for condensates before and after recruitment of peptides. Black line indicates the mean. Bottom: Nanostars were designed to be sequence-orthogonal, thus their one-pot assembly with fluorescently labeled clients shows their recruitment and compartmentalized into specific condensates (red, blue, and yellow). (C) Sequence-orthogonal single-stranded RNA motifs pairs (two nanostar systems, 3sβ and 3sγ), modified to include Corn and Red Broccoli fluorogenic aptamers. To reduce steric hindrance, 25% of 3sβ1 or 3sγ1 were modified to include FLAPs, while the other 75% and all 3sβ2 or 3sγ2 remained unmodified. The peaks of the FITC/Cy3 angle histogram for individually annealed condensates (single-color) align with the peaks of the one-pot assembly (multi-color). Experiments were done in triplicates; images provide representative examples. Violin plots and histograms pool data from all replicates. Scale bars: 20 μm.

Next, we show that our RNA condensates can recruit client peptides to the dense phase. For this purpose, we replaced FLAPs with peptide and protein binding domains: we considered a streptavidin binding aptamer (appended to nanostar variant 4m3), boxB RNA that binds P22 N peptide (appended to nanostar variant 4m5), and TAR RNA that binds TAT peptide (appended to nanostar variant 4m6)^39–41^, shown in Fig. 3B. We selected these peptides for their versatility: the streptavidin-biotin pair is widely used for purification and localization assays^42^; the P22 bacteriophage N peptide is a strong binder for its aptamer boxB, and has been useful for RNA colocalization studies^43^; finally, Tat bound to its aptamer is a strong transcriptional regulator involved in HIV replication^44^. Strands including aptamers were doped into the mixture of unmodified strands at a ratio of 1:4 to prevent steric hindrance of sticky-end hybridization. We verified that each peptide binding nanostar yields condensates when assembled individually, and it successfully recruits its client; after peptide recruitment, condensates become on average larger (Fig. 3B, bottom). Undesired cross-binding of targets to their non-cognate aptamer occurs only for TAT, due to its positively charged amino acid residues that causes non-specific binding to negatively charged RNA (Fig. S7). Overall, our orthogonal RNA nanostars partition the fluorescently labeled peptides into distinct compartments that do not mix.

Finally, we confirmed that also single-stranded RNA nanostars can recruit client molecules. We selected orthogonal two-nanostar variants 3sβ and 3sγ and we modified them to include FLAPs binding DFHO (Red Broccoli and Corn, respectively) as an additional arm. We observed the formation of condensates of distinct colors upon annealing nanostars together (Fig. 4C, Fig. S15). We verified that these condensates do not mix through our quantitative image analysis (computation of the arctangent of FITC/Cy3 intensity ratio for each pixel, Fig. S14), which shows aligned histogram peaks when purified nanostars are annealed separately or together.

### Cotranscriptional formation of RNA condensates

We finally demonstrate that single-stranded RNA nanostars produce condensates cotranscriptionally at 37°C, in the absence of thermal annealing (Fig. 5A). Tandem stem-loops fold based on local RNA interactions, so they are expected to form stable structures as they are being transcribed under isothermal conditions^45^. We transcribed nanostars using linear templates under the control of the T7 bacteriophage promoter, using buffer for high-yield *in vitro* transcription that includes MgCl_2_ ^46^ (see Methods). We observed large condensates (stained with SYBR Gold) within 1-2 hours of transcription of nanostar variant 3sv2-stem1 (Fig. 5A), which was selected from our library as it produced abundant spherical condensates when purified and thermally treated with a 40°C hold as well as in the presence of MgCl_2_ (Fig. 3F, Fig. S11). The average condensate size doubles within the first 3 hours as confirmed by the mean-chord length (μ_CLD_) bar plot (Fig. 5B); larger aggregates can be obtained by increasing the DNA template concentration (Fig. S16). The number of condensates only slightly decreases over time, which suggests that our reaction conditions promote condensate growth primarily by monomer addition, rather than fusion, while nanostars are being transcribed. We also tested the cotranscriptional assembly of variant 3sβ, which similarly produces aggregates that grow rapidly into a slowly fusing network (Fig. 5C-D). We found that the amount of monovalent and divalent cations (NaCl and MgCl_2_) in the transcription buffer has significant effects on cotranscriptional formation, as shown in control experiments involving the two-nanostar variant 3sβ (Fig. S17); buffers included in commercial kits may fail to yield condensates if they do not include sufficient cation levels. We expect that cotranscriptional condensates can be obtained using other nanostar variants that produce condensates from purified RNA with the melt and 40°C hold protocol (Fig. 3F, Fig. S10).

**Figure 5:**
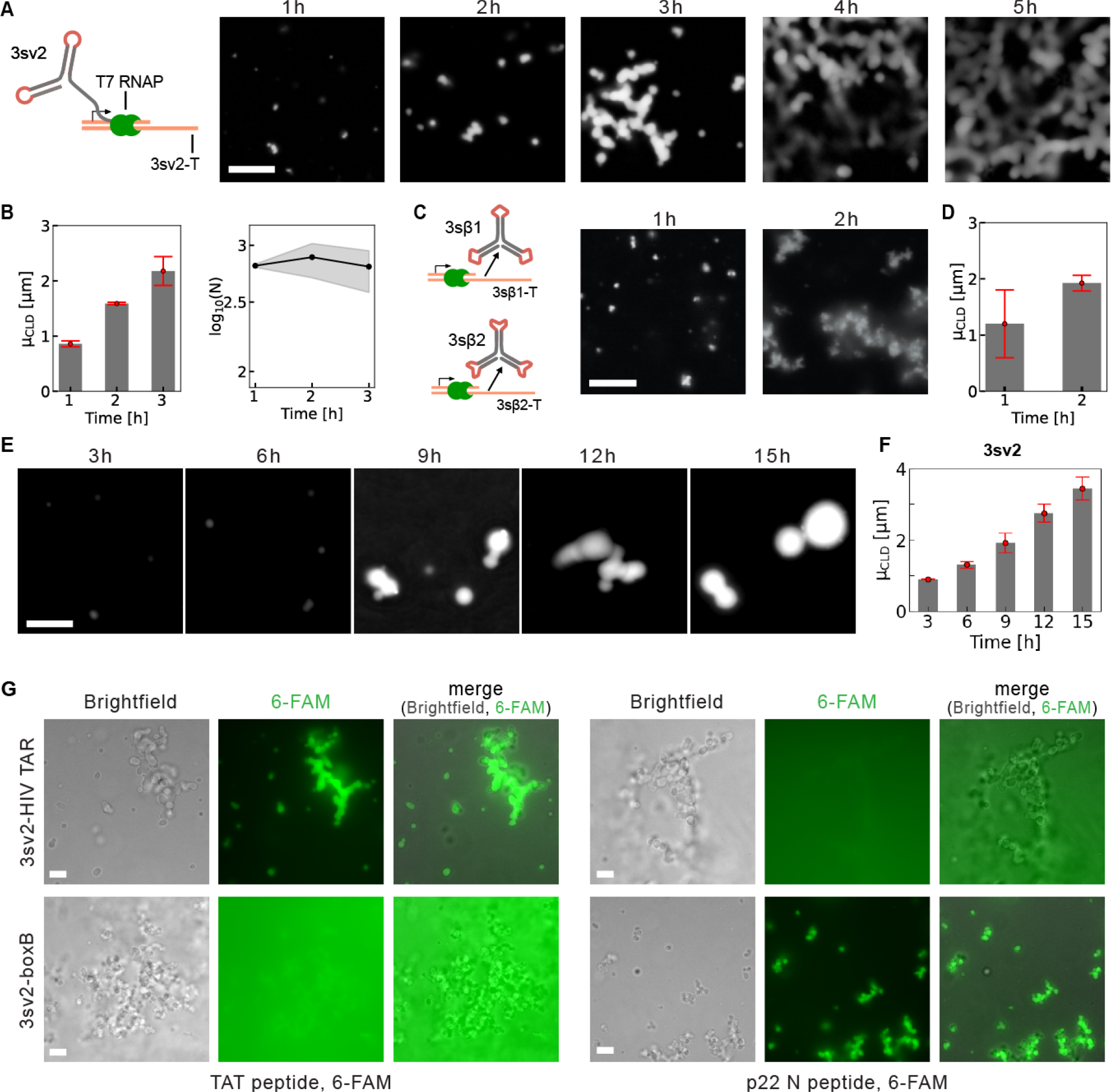
Cotranscriptional, isothermal formation of single-stranded RNA nanostars and peptide recruitment. (A) Scheme shows transcription and cotranscriptional folding of motif 3sv2; example images show condensate formation in T7 in-vitro transcription reaction sampled at different time points. (**B**) Temporal evolution of the mean-chord length (μ_CLD_, left) and number (N, right) of 3sv2 nanostars. (**C**) Scheme shows transcription and cotranscriptional folding of motif pairs 3sβ1 and 3sβ2; example images show 3sβ condensate growth in T7 in-vitro transcription reaction. (**D**) Temporal evolution of μ_CLD_ of 3sβ condensates; CLD analysis is not meaningful after 2-3 hours of incubation due to the growth of the RNA dense phase. (**E**) Example images show 3sv2 condensate growth in a cell-free PURExpress® reaction. (**F**) Temporal evolution of μ_CLD_ of 3sv2 condensate in the cell-free system. B, D and F show mean (bar heights or solid line) ± standard error (red error bar or shaded area) from eleven FOVs within three replicates. (**G**) Brightfield and fluorescent images show specific peptide recruitment to condensates formed cotranscriptionally 2 hours after incubation. 3sv2 nanostars were modified to include TAR (top) or boxB (bottom) aptamer, recruiting either TAT or p22 N peptide labeled with 6-FAM. Peptides are added before imaging. Experiments were done in triplicates; images are representative examples. Bar charts show the mean of three replicates; brackets and shaded areas represent the standard deviation of the mean over three replicates. Scale bars: 10 μm.

Cotranscriptionally produced RNA condensates could be useful as membraneless organelles that can spontaneously recruit proteins. To demonstrate that this is immediately feasible in cell-free systems, we verified that nanostar 3sv2-stem1 could produce condensates when transcribed using PURExpress®, a commercial kit for cell-free protein synthesis^47^. The production of large condensates appears slow in PURExpress® when compared to a high-yield transcription kit, as evidenced by example images of Fig. 5E and by the μ_CLD_ shown in Fig. 5F. Condensate growth rate may be increased by using larger amounts of DNA template or of RNA polymerase. Finally, we modified 3sv2-stem1 to recruit p22 N and TAT peptides using their corresponding aptamers (boxB and TAR, respectively) and demonstrated that these peptides are recruited to the condensates cotranscriptionally. After incubating DNA templates for 2 hours in a high-yield transcription buffer, we added peptides to our samples, verifying the correct, specific recruitment of peptides to the dense phase (Fig. 5G).

## Discussion

We have demonstrated the design and synthesis of modular RNA nanostars for phase separation. We showed that both multi-stranded and single-stranded nanostars robustly form condensates in standard buffers commonly adopted in nanotechnology applications, in the absence of binding partners. We focused on two protocols, (1) transcribing and purifying RNA strands, then performing a temperature treatment consisting of a denaturing step and a long temperature hold, or (2) cotranscriptional assembly. We examined how condensation is influenced by various nanostar design features and extrinsic factors such as ionic conditions and temperature. Further, we have programmed orthogonal condensates to recruit and organize small molecules and peptides, thus mimicking the ability of biological condensates to recruit clients^48^. Our strategy is modular, and may lead to the development of libraries of orthogonal condensates recruiting diverse clients. Multi-stranded and single-stranded RNA nanostars offer different advantages depending on the downstream application or purpose. Owing to the ease in designing linear sticky-ends and hybridizing domains, the affinity and valency of multi-stranded nanostars can be easily tuned, making it possible to build RNA condensates with a broad range of biophysical properties that should be comparable to those demonstrated for similar DNA condensates^19,20,27^. Further, this design allows for modular introduction of RNA or DNA strands with distinct functionalities, including client recruitment and adaptation to chemical or physical inputs that are relevant for developing therapeutic, multifunctional biomaterials^49^. Single-stranded nanostars produce condensates isothermally under physiological conditions, making them immediately useful as RNA organelles that can be produced in artificial cells, as demonstrated in a parallel study^25^. We are evaluating whether these constructs can be genetically encoded in living cells.

A distinguishing feature of our RNA condensates is that they are formed through compact, nanostructured motifs designed using short sequences that are optimized to fold as desired. While RNA coacervates can emerge from unstructured RNA homopolymers, their formation requires the presence of polymeric cations that are not necessary in our system^50–52^. Previously, RNA strands assembling into pure RNA hydrogels have been identified via SELEX^53^, however, this approach does not offer a clear design strategy to modularly adapt the condensing motifs as easily as with RNA nanostars. Recent work demonstrated that liquid RNA droplets can be produced inside cells through long RNA molecules featuring short sequence repeats (CAG/CUG)^15,16,54^. By fusing the CAG-repeats to recruitment domains, it was possible to demonstrate compositional control of RNA droplets both in bacteria and in mammalian cells^15,16^. Unlike RNA nanostars, these long repeats are expected to form tangled complexes that require annealing to form gels *in vitro*, and rely on energy-dissipating cellular machinery to remain in a liquid state^15^. Intracellular RNA condensates were also obtained via homodimerization of two different RNA aptamer repeats^55^, which remain demixed forming up to two types of orthogonal compartments inside cells. In these exciting achievements, sequence repeats are an expedient strategy to introduce multivalency; however, it is not clear whether these methods can be used to build a scalable number of orthogonal condensates.

A simplified working model of RNA nanostars is that we can determine their affinity, valency, and size by changing the nanostar sticky-ends/kissing loops, number of arms, and arm length. However, we still lack a clear picture of how different combinations of these parameters affect phase transitions and condensation kinetics. Further, some of these parameters mutually couple in ways that are difficult to predict; for example, changes in the stem sequence can have an impact on the nanostar affinity if they affect the folding of the interaction domains, and these effects are sensitive to temperature and ionic conditions. The influence of combinations of design changes could be predicted by coarse-grained models that capture the interactions among nanostars^56,57^, and should be validated against systematic experiments characterizing phase diagrams and growth rates of nanostar condensates. These models could accelerate the design of customized RNA condensates while minimizing experimental burden. Finally, while we have demonstrated modular recruitment of small molecules and peptides, we expect limitations in which classes of molecules of proteins are recruitable by RNA nanostars with specificity. RNA aptamers can be promiscuous, for example TAR RNA is known to bind positively charged arginine^58^. Further, arbitrary RNA domains may non-specifically recruit amino acid sequences; for example, positively charged peptides naturally tend to bind to the negatively charged backbone of RNA, regardless of sequence specificity (like we observed with TAT protein). We expect that, in complex biological samples, various RNA-binding proteins could bind non-specifically to our artificial condensates and possibly compromise their formation. Another challenge is that some recruitment motifs could introduce misfolding or remodeling of the secondary structure of contranscriptionally produced single stranded nanostars. Design methods developed for RNA origami may provide valuable lessons toward enhancing the robustness and the functionalities of RNA nanostars^59^.

Orthogonal, customizable micron-sized compartments made with RNA may find clinical and industrial applications for purification, diagnostics, and therapeutic treatments. RNA condensates with the capacity to selectively sequester binding targets may be useful for separating crude mixtures of molecules or low-volume clinical samples. The incorporation of stimuli-responsive domains in RNA nanostars could make it possible to develop sensors and diagnostic tools, and further engineering may allow this platform to serve as a smart drug delivery system capable of encapsulating and releasing therapeutic molecules. Lastly, we envision that single-stranded RNA nanostars may be used to produce functional RNA organelles in living cells and potentially control cellular processes.

## Methods

Sequences, detailed methods, and additional experiments are included in the Supplementary Table and Supplementary Information file.

### Sequence design

The RNA strands for the multi-stranded motifs (4m) were designed and optimized using the NUPACK design tool using scripts reported in SI Section 1^26^. Aptamer sequences were selected from previous studies^37,38^. Sequences were appended to the 5’-sticky end of the S1 strand of a given motif. The secondary structure of the motif with aptamer modification was analyzed using NUPACK to ensure that the secondary structure of the motif and aptamer was similar to the fold of the individual modules. Three-armed single-stranded RNA nanostars (3s) were designed by combining distinct arm sequences with several kissing loop variants. Stem 1 sequences were adapted from the multi-stranded motifs (4m); stem 2 sequences were adapted from the design by Sato et al. for DNA nanostars^17^; stem 3 sequences were designed using NUPACK. Two of the three spacers between arms include two unpaired adenines, and the other is a nick, allowing the motif to have flexible configurations. All KLs are nine nucleotides long and include a six-nt interaction sequence flanked by three unpaired adenine residues, two upstream and one downstream of the interaction sequence (5’-AA…A-3’). KL variants used for the single-nanostar motifs were adapted from the HIV-1 palindromic KL sequence^32^. Each variant was obtained by introducing a single-base substitution in the six nt interaction domain of the wild type KL (5’-GCGCGC). KL for two-nanostar motifs were designed *de novo* to function as pairs (heterodimers). All sequences include four GC pairs and two AU pairs to ensure similar bond stabilities. Fluorogenic aptamer sequences were adopted from literature ^37,38^ and modified using the same method adopted for multi-stranded designs. Expected folding of each design was confirmed using NUPACK ^26^. All sequences are listed in the Supplementary Table.

### RNA synthesis

RNA strands were transcribed from PAGE purified DNA templates, including a T7 promoter, purchased from Integrated DNA Technologies. Lyophilized DNA was resuspended in nuclease-free water and DNA template was annealed in 1X TE/50 mM NaCl from 90°C to RT at -1°C/min. RNA strands were individually transcribed in vitro using the AmpliScribe T7-Flash transcription (ASF3507, Biosearch Technologies) kit from DNA templates. RNA strands were then purified using Amicon Ultra 10K 0.5 ml centrifugal filters and 1X TE buffer and centrifuging three times at 14,000 g. For multi-stranded nanostars each strand was transcribed from a fully double stranded DNA template including the T7 promoter. For single-stranded nanostars we used single stranded non-coding DNA templates annealed to a 21-nt complement including the promoter region and a 4 nt sealing domain.

### Condensate preparation

Condensates were formed in assembly buffer including 40 mM HEPES, 100 mM KCl, 500 mM NaCl. Strands were thermally annealed in an Eppendorf Mastercycler using a melt and hold protocol which includes a melt at 70 °C for 10 minutes, followed by 12 hours of incubation at specified temperatures, and by a quick drop to 20 °C for 5 minutes before imaging. For multi-stranded nanostars, we used purified RNA strands each at equimolar concentrations (5 μM). Strands including aptamer sequences were added at 1.25 μM concentration (25% doping). For single-stranded nanostars we used purified RNA with a concentration of 5 μM. For the two-nanostar condensates we used purified RNA with each nanostar at 5 μM concentration; thus, the total RNA concentration in these experiments was doubled. For two-nanostar motif experiments using fluorogenic RNA aptamers, aptamers-containing strands were added at a 25% doping.

### Preparation of peptide and protein targets

Fluorescent peptides were synthesized by either LifeTein, LLC or GenScript. Fluorophores were added to the N-terminal, AlexaFluor647 for P22 N peptide, and 6-FAM for TAT peptide. For cotranscriptional experiments, both N peptide and TAT were labeled with 6-FAM. Peptide synthesis was guaranteed a purity of ≥95% with standard Trifluoroacetic acid (TFA) removal (Final TFA Counterion % < 10%. Lyophilized peptide was resuspended in 10 mM HEPES with a molarity of ∼350 μM and stored at 5°C. Streptavidin, Alexa Fluor™ 555 conjugate was purchased from Thermo Fisher Scientific at 2 mg/ml concentration, with a molarity of ∼35 μM. Streptavidin conjugate was diluted with 10 mM HEPES and stored at 5°C.

### Cotranscriptional production of condensates

DNA templates were prepared as described under RNA synthesis. For the single-stranded RNA nanostars, the RNA strands were transcribed *in vitro* at 37°C using 7.5% (v/v) T7 polymerase from the AmpliScribe T7-Flash transcription kit (ASF3507, Biosearch Technologies), 40mM of Tris-HCl, 10 mM of NaCl, 30 mM MgCl_2_, 2 mM spermidine, 7.5 mM each NTP, 10 mM DTT. For the sequence-orthogonal single-stranded RNA nanostars (two-nanostar system), the RNA strands were transcribed *in vitro* under the same conditions, with the exception of the NaCl and MgCl2 concentrations, which were adjusted to 20 mM each. When using the PURExpress® kit (E6800S, New England Biolabs), we adhered to the manufacturer’s protocol and incubated our sample at 37°C. Unless otherwise specified, we used a final concentration of 10 nM DNA template for the *in vitro* transcription; we used 10 ng DNA for the cell-free PURExpress® reaction.

### Fluorescence microscopy

Condensates produced from multi-stranded motifs were obtained with an Olympus BX-UCB upright fluorescence microscope using a 20x air objective. We used filtersets Cy3 (Chroma Filter Set Exciter D540/25x EX Dichroic Q565lp BS Emitter D620/60m EM), Cy5 (Chroma Filter Set Exciter HQ620/60x EX Dichroic Q660LP BS Emitter HQ700/75m EM), and FITC (Chroma Filter Set Exciter D480/30x EX Dichroic Q505lp BS Emitter D535/40m EM), with a standard exposure time of 100 ms for samples stained with SYBR gold, 500 ms for samples with DFHO or fluorescent peptides and proteins. Condensates produced from single-stranded motifs were imaged using a Nikon Eclipse TI-E inverted microscope using a 60x oil immersion objective. SYBR gold-stained samples were detected using the FITC channel (ex 455 - 485 nm/em 510 - 545 nm) with an exposure time of 100 ms. DFHO-stained samples were detected in both the FITC channel with an exposure time of 200 ms, and in the Cy3 channel (ex 512 - 552 nm/em 565 - 615 nm) with an exposure time of 100 ms. For samples stained with SYBR Gold, 1X SYBR gold was used after thermal treatment. For samples with DFHO, 50 μM DFHO in 2mM HEPES buffer was used before thermal treatment (for multi-stranded nanostars) and after thermal treatment (for single-stranded nanostars).

### Image processing and visualization

All fluorescence images were processed in FIJI (ImageJ). Raw images were background subtracted, contrast enhanced, and converted to a binary mask as described in detail in the SI. For condensate number analysis, objects smaller than 6 px^2^ were considered noise and excluded. Condensate numbers were autogenerated by FIJI and recorded. To gather information on condensate size, we measured chord length distributions (CLD)^28–31^ from the binary masks using a Python3 script based on PoreSpy, which relies on Scipy and Skimage. Violin plots were generated using a Python3 script based on Seaborn. All experimental replicates were pooled to a single violin plot. Means were computed across three technical replicates. To determine whether our nanostars including distinct fluorogenic aptamers produce condensates that mix or do not mix, we built pixel intensity histograms from fluorescence microscopy images (Fig. 4, SI Fig. S15). This was done because upon DFHO staining, both Corn and Red Broccoli aptamers can be detected in the FITC channel and the Cy3 channel, albeit with varying intensity, since the Corn aptamer emission peak is 545 nm, and the Red Broccoli emission peak is 582 nm. Histograms were generated from the coordinate angles calculated from every pixel within the regions of interest. Additional details on image processing and visualization are provided in the SI.

## Acknowledgements

JMS is a Merck Awardee of the Life Sciences Research Foundation. EF acknowledges support from the US NSF through CAREER award 1938194 and FMRG: Bio award 2134772, and from the Sloan Foundation through award G-2021-16831. LDM acknowledges support from the European Research Council (ERC) under the Horizon 2020 Research and Innovation Programme (ERC-STG No 851667 – NANOCELL) and a Royal Society University Research Fellowship (UF160152, URF\R\221009). GF acknowledges funding from the Department of Chemistry at Imperial College London.

