## Supplementary Information for "Modular RNA motifs for orthogonal phase separated compartments"

**Table of Contents**

1 Materials and Methods 2
1.1 Sequence design 2
1.1.1 Multi-stranded nanostars (Caltech) 2
1.1.2 Three-arm single-stranded nanostars (UCLA) 4
1.2 RNA synthesis 4
1.2.1 Multi-stranded nanostars (Caltech) 4
1.2.2 Three-arm single-stranded nanostars (UCLA) 4
1.3 Sample Preparation 5
1.3.1 Multi-stranded nanostars (Caltech) 5
1.3.2 Three-arm single-stranded nanostars (UCLA) 6
1.4 Imaging 6
1.4.1 Multi-stranded nanostars (Caltech) 7
1.4.2 Three-arm single-stranded nanostars (UCLA) 7
2 Supplementary Experiments and Text 10
2.1 Multi-stranded nanostars (Caltech) 10
2.2 Three-arm single-stranded nanostars (UCLA) 14
References cited 21

### 1 Materials and Methods

#### 1.1 Sequence design

##### 1.1.1 Multi-stranded nanostars (Caltech)

**Multi-stranded RNA Nanostar Design** The RNA strands for the multi-stranded motifs (4m) were designed and optimized using the NUPACK Design tool with the script shown below[1]. A target secondary structure of the multi-stranded motif was denoted by DU+ notation. Sticky ends were pre-selected and noted as a design restraint as well as certain repeat nucleotide sequences. We used the default energy parameters from Serra and Turner, 1995 in 1M Na+ [1]. Aptamer sequences were selected from previous studies [2,3]. Sequences were appended to the 5'- sticky end of the S1 strand of a given motif. The secondary structure of the motif with aptamer modification was analyzed using NUPACK to ensure that the secondary structure of the motif and aptamer was similar to the fold of the individual modules.

```
---------------------------------------------------------------------------------------------------------
#
#NUPACK script for multi-stranded motif strand design
#
material = rna
temperature[C] = 25.0    # optional units: C (default) or K
trials = 3
sodium[M] = 1.0    # optional units: M (default), mM, uM, nM, pM
dangles = some

#
# target structure using DU+ notation
#


#5' SE
structure msmotif = U7 D15 (U2 D15 (+ U7) U2 D15 (+ U7) U2 D15 (+ U7) U2)

# sequence domains
#
domain A2 = AA

#input sticky-end sequence
domain SE = GCUAGCA
domain S4 = N15
domain S3 = N15
domain S2 = N15
domain S1 = N15
```

#
### thread sequence domains onto target structures
#

#5' SE
msmotif.seq = SE S1 A2 S2 SE S2* A2 S3 SE S3* A2 S4 SE S4* A2 S1*

#
### specify stop conditions for normalized ensemble defect
### default: 1.0 (percent) for each target structure
#

msmotif.stop = 1.0

#
### prevent sequence patterns
#
prevent = AAAA, CCCC, GGGG, UUUU, KKKKKK, MMMMMM, RRRRRR, SSSSSS, WWWWWW, YYYYYY
#
---------------------------------------------------------------------------------------------------------

**Aptamer design**

Fluorogenic aptamers: The Corn aptamer was modified from previous work [3] by adding GG to the 5' end and CC to the 3' end to ensure transcription. Red Broccoli and Orange Broccoli aptamers were modified from [2] to include flanking sequences GUAUGUGG at the 5' end and CCCACAUAC at the 3' end, which surround conserved fluorogen-binding sequences and broccoli bridging sequences.

Corn:
Underlined bases were added to the original sequence found in [3]
5'- GGCGCGAGGAAGGAGGUCUGAGGAGGUCACUGCGCC -3'

Red Broccoli:
flanking region- **fluorogen binding region-conserved-** broccoli region-*RB tuning base*-**fluorogen binding region-conserved-**flanking region
5'-GUAUGUGG-**GAGACGGUCGGGUCC-**AGAUAUUCGUAUCUGUCGAGUAG-*U*-**GUGUGGG CUC-**CCACAUAC-3'

Orange Broccoli:
flanking region- **fluorogen binding region-conserved-** broccoli region-*OB tuning base*-**fluorogen binding region-conserved-**flanking region
5'-GUAUGUGG-**GAGACGGUCGGGUCC-**AGAUAUUCGUAUCUGUCGAGUAG-*C*-**GUGUGGG CU-**CCCACAUAC-3'

Peptide and protein binding aptamers: The boxB RNA sequence that binds P22 N peptide and HIV-1 Tat aptamer that binds Tat peptide was taken from previous work[4,5]. The streptavidin binding aptamer was modified from previous work[6] by adding a G at the 5' end and a C at the 3' end to ensure transcription.

**boxB:** 5'-GGUGCGCUGACAAAGCGCGCC-3'

**HIV-1 Tat:** 5’-GGAGCUUGAUCCCGGAAACGGUCGAUCGCUCC-3’
**Streptavidin:**
5’-GAUGCGGCCGCCGACCAGAAUCAUGCAAGUGCGUAAGAUAGUCGCGGGUCGGCGGCCGCAUC-3’
Underlined bases were added to the original sequence

##### 1.1.2 Three-arm single-stranded nanostars (UCLA)

Three-armed single-stranded RNA nanostars (3s) were designed by combining two distinct arm sequences with several kissing loop variants.
Stem 1 arm sequences were designed by adapting the sequences of the multi-stranded motif (15 bp arms, see sequence section). Stem 2 arm sequences was adapted from the motif designed for three-arm DNA nanostars by Sato et al. (16 bp arms,[7]). Stem 3 arm sequences were designed using Nupack (20 bp arms). Two of the three spacers between arms include two unpaired adenines, and the other is a nick, allowing the motif to have flexible configurations.
We tested two sets of kissing-loop (KL) sequences. All KLs are nine nucleotides long and include a six-nt interaction sequence flanked by three unpaired adenine residues, two upstream and one downstream of the interaction sequence (5’-AA…A-3’). The first set of KLs (single nanostar motif) was adopted from the HIV-1 palindromic KL sequence following previous work [8]. Each variant was obtained by introducing a single-base substitution in the six nt interaction domain of the wild type KL (5’-GCGCGC) . The second set (two nanostar motifs) features KL sequences designed *de novo* to function as pairs (heterodimers). This eliminates the restriction of using a palindromic kissing loop sequence and expands the possible designs. All sequences include four GC pairs and two AU pairs to ensure similar bond stabilities.
The fluorogenic aptamer sequences were adopted from the literature[2,3] and modified using the same method adopted for multi-stranded designs (Section 1.1.1).
Minimum free energy configurations of each design were confirmed using NUPACK[1].

---

#### 1.2 RNA synthesis

---

##### 1.2.1 Multi-stranded nanostars (Caltech)

RNA strands were transcribed from PAGE purified DNA templates, including a T7 promoter, purchased from Integrated DNA Technologies. Lyophilized DNA was resuspended in nuclease-free water and DNA template was annealed in 1X TE/50 mM NaCl from 90°C to RT at -1°C/min. RNA strands were individually transcribed *in vitro* using the AmpliScribe T7-Flash transcription (ASF3507, Biosearch Technologies) kit from DNA templates. RNA strands were then purified using Amicon Ultra 10K 0.5 ml centrifugal filters and 1X TE buffer and centrifuging three times at 14,000 g. Concentrations of DNA and RNA were determined by absorption measurements using a Nanodrop 2000c.

##### 1.2.2 Three-arm single-stranded nanostars (UCLA)

RNA strands were transcribed from partially single-stranded DNA templates containing a T7 promoter. A “sealing domain” (5’-GCGC-3’) was added upstream of the T7 promoter sequence (5’-TAATACGACTCACTATA-3’). The non-coding strands of the DNA templates were purchased from Integrated DNA Technologies (IDT). The non-coding strands were annealed to the 21 nt complement of the T7 promoter.

For *in vitro* studies using purified RNA, DNA templates with concentrations of 2 μM were annealed with the complementary promoter strand in 1X TE/50 mM NaCl by holding at 90°C for 5 min to allow denaturing, then cooling to RT at -1°C/min. RNA strands were individually transcribed *in vitro* using the AmpliScribe T7-Flash transcription kit (ASF3507, Biosearch Technologies) from 200 nM DNA templates at 37°C for 4 hours, and then treated with DNase for 15 minutes at 37 °C. RNA strands were then purified using Amicon Ultra 10K 0.5 ml centrifugal filters (cat# UFC501096) and 1X TE buffer and centrifuging three times for 20 minutes at 14,000 g at 20 °C. We then calculated the concentration of the purified products based on the absorption at 260 nm (A260) with Nanodrop 2000c and the extinction coefficient provided by the manufacturer. The purified RNA was then resuspended to 50 μM stock in 1x TE buffer and stored at -80°C for later use.
For co-transcription studies, the DNA templates were prepared as previously described. For the single-stranded RNA nanostars, the RNA strands were transcribed *in vitro* at 37°C using 7.5% (v/v) T7 polymerase from the AmpliScribe T7-Flash transcription kit (ASF3507, Biosearch Technologies), 40mM of Tris-HCl, 10 mM of NaCl, 30 mM $MgCl_2$, 2 mM spermidine, 7.5 mM each NTP, 10 mM DTT. For the sequence-orthogonal single-stranded RNA nanostars, the RNA strands were transcribed *in vitro* under the same conditions, with the exception of the NaCl and MgCl2 concentrations, which were adjusted to 20 mM each. When using the PURExpress® kit, we adhered to the manufacturer's protocol and incubated our sample at 37°C. Unless otherwise specified, a final concentration of 10 nM DNA template was used for the *in vitro* transcription, and 10 ng DNA was used for the cell-free reaction.

#### 1.3 Sample Preparation

##### 1.3.1 Multi-stranded nanostars (Caltech)

**Preparation of condensates** Condensates formed by multi-stranded motifs were prepared with purified RNA strands at equimolar concentrations (5 μM) in our assembly buffer including 40 mM HEPES, 100 mM KCl, 500 mM NaCl. Strands were thermally annealed in an Eppendorf Mastercycler using the melt and hold protocol discussed in the manuscript (Fig. 2B in the manuscript), which includes a melt at 70 °C for 10 minutes, followed by 12 hours of incubation at specified temperatures, and by a quick drop to 20 °C for 5 minutes before imaging. Strands including aptamer sequences were added at 1.25 μM concentration (25% doping).

**Preparation of peptide and protein targets** Fluorescent peptides were synthesized by either LifeTein, LLC or GenScript. Fluorophores were added to the N-terminal, AlexaFluor647 for P22 N peptide, and 6-FAM for TAT peptide. Peptide synthesis was guaranteed a purity of ≥95% with standard Trifluoroacetic acid (TFA) removal (Final TFA Counterion % < 10%. Lyophilized peptide was resuspended in 10 mM HEPES with a molarity of ~350 μM and stored at 5°C. Streptavidin, Alexa Fluor™ 555 conjugate was purchased from Thermo Fisher Scientific at 2 mg/ml concentration, with a molarity of ~35 μM. Streptavidin conjugate was diluted with 10 mM HEPES and stored at 5°C.
Peptides were added to the condensate samples after the melt and hold step.

**Peptide sequences**
P22 N peptide: GNAKTRRHERRRKLAIERDTIGY
HIV-1 TAT: SFITKALGISYGRKKRRQRRRPPQGSQTHQVSLSKQ

##### 1.3.2 Three-arm single-stranded nanostars (UCLA)

**Melt and hold condensate preparation** Condensates formed by single-stranded motifs were prepared in assembly buffer including 40 mM HEPES, 100 mM KCl, and 500 mM NaCl. To form single-nanostar condensates with palindromic KLs, we used purified RNA with a concentration of 5 µM. For two-nanostar condensates, we used purified RNA with each nanostar at 5 µM concentration; thus, the total RNA concentration in these experiments was doubled. For two-nanostar motif experiments using fluorogenic RNA aptamers, aptamers-containing strands were added at a 25% doping ratio (thus, the final purified RNA concentrations in the annealing mix were 3.75 µM non-modified strand and 1.25 µM aptamer-including strand). We annealed the strands using the melt and hold protocol (melt at 70 °C for 10 minutes, followed by 12 hours of incubation at specified temperatures, and by a quick drop to 20 °C for 5 minutes before imaging). We expect that condensates start forming during the temperature drop from 70 to 50 °C, grow during the 12-hour incubation period, and become “frozen” when the temperature decreases to 20 °C.

Condensates formed from motifs without aptamer were imaged with 1X SYBR Golding staining (S11494, Thermofisher). Dye was added 5 minutes after cooling to 20°C. Condensates formed from aptamer-appended motifs were stained with DFHO (Lucerna Technologies). DFHO stock was stored in DMSO at a concentration of 10 mM. DFHO staining solution was prepared by diluting DFHO stock in HEPES buffer with a final concentration of 1 mM DFHO and 40 mM HEPES. Samples were imaged immediately after staining.

**Cotranscriptional condensate preparation** Cotranscriptional experiments were conducted as previously described in section 1.2.2. RNA strands were transcribed in vitro at 37°C with 1.5 uL of T7 polymerase from the AmpliScribe T7-Flash transcription kit (ASF3507, Biosearch Technologies), 40mM of Tris-HCl, 10 mM of NaCl, 30 mM $MgCl_2$, 2 mM spermidine, 7.5 mM each NTP, 10 mM DTT.

When using the PURExpress® kit (E6800S, New England Biolabs), we adhered to the manufacturer’s protocol for buffer assembly and we incubated our samples at 37°C.

Unless otherwise specified, a final concentration of 10 nM DNA template was used.

**Preparation of peptide targets and cotranscriptional peptide recruitment experiments** Peptides were purchased from GenScript. Both P22 N and TAT peptide were labeled using 6-FAM at the N-terminal. Peptide synthesis was guaranteed a purity of ≥95% with standard Trifluoroacetic acid (TFA) removal (Final TFA Counterion % < 10%). and resuspended in water to a 10x concentration (10 µM) and used at 1X (1 µM). DNA templates (10 nM) for nanostars carrying the peptide aptamer were transcribed for 2 hours at 37°C before adding their corresponding peptides (1 µM final). To test peptide binding specificity towards their associated aptamer, a cross experiment was run at the same time, adding P22 N peptide to TAT nanostars and TAT-peptide to boxB nanostars (n=3 independant transcriptions for both normal and cross experiments). For each replicate, we collected between 5 to 13 images showing a representative sample of the condensate morphology and fluorescence intensity. Both green fluorescent channel and brightfield were captured, to show presence of condensates in case of an absence of recruitment of the fluorescent peptides. Triplicates were consistent, and a specific recruitment of the peptides was observed for both boxB / N peptide and TAT / TAT-peptide pairs.

**Peptide sequences** (same as in the multi stranded nanostar experiments)
P22 N peptide: GNAKTRRHERRRKLAIERDTIGY
HIV-1 TAT: SFITKALGISYGRKKRRQRRRPPQGSQTHQVSLSKQ

#### 1.4 Imaging

##### 1.4.1 Multi-stranded nanostars (Caltech)

**Imaging** Images of multi-stranded motifs were obtained with an Olympus BX-UCB upright fluorescence microscope. Specified filtersets Cy3 (Chroma Filter Set Exciter D540/25x EX Dichroic Q565lp BS Emitter D620/60m EM), Cy5 (Chroma Filter Set Exciter HQ620/60x EX Dichroic Q660LP BS Emitter HQ700/75m EM), and FITC (Chroma Filter Set Exciter D480/30x EX Dichroic Q505lp BS Emitter D535/40m EM) were used to image condensates with a 20x air immersion objective used to collect all images, with a standard exposure time of 100 ms for samples stained with SYBR gold, 500 ms for samples with DFHO or fluorescent peptides and proteins. For samples stained with SYBR Gold, 1X SYBR gold was added after thermal treatment. For samples with DFHO, 50 µM DFHO was added before thermal treatment. Before imaging, coverslips were rinsed with acetone, isopropyl alcohol, then with milliQ water and dried with a kim wipe.

**Image enhancement** All fluorescence images in figures were processed in FIJI (ImageJ) with a background subtraction with a rolling ball radius of 25-50 pixels.

**Image analysis** For peptide recruitment samples, size analysis of each condition was performed using data from three experimental replicates and FOV=10.
Masks were generated from the CY3 channel for the Streptavidin-binding motif, the CY5 channel for the P22-binding motif, and FITC channel images for TAT-binding motifs. Images were thresholded using a custom macro implementing the following pipeline: contrast enhancement (5% pixels saturated), background subtraction with white top-hat morphological filters (element=disk, radius=20), denoising via Gaussian Blur (sigma=2), and auto threshold using the MaxEntropy method.
For incubation over time samples (Fig. 2D), size analysis of each time point was performed using data from one experimental replicate and FOV=14.
Images were threshold using a custom macro implementing the following pipeline: contrast enhancement (5% pixels saturated), denoising via Gaussian Blur (sigma=2), and background subtraction with rolling ball method (radius = 0.5) for five times, and auto thresholded using the MaxEntropy method. Masks were saved as TIFF files for droplet size analysis.
Droplet size information was extracted as Chord Length Distribution (CLD) from the binary masks using a Python3 script based on PoreSpy, particularly Porespy.filters.apply_chords, which relies on Scipy and Skimage. The selected options allow for adjacent chords (spacing = 1) and include objects touching the edges of the FOV (trim_edges = False).
For each binary mask, the script calculates chords along the x and y directions via the chord_counts function from PoreSpy. The resulting Numpy arrays are saved in NPY format via Numpy. Each sample's resulting chord lengths, along x and y and across all FOVs, are pooled together, and saved into new arrays. The reported mean are calculated as mean(mean_sample1, mean_sample2, mean_sample3).

##### 1.4.2 Three-arm single-stranded nanostars (UCLA)

**Imaging** Fluorescence microscopy images were obtained with an inverted microscope (Nikon Eclipse TI-E) with a 60x oil immersion objective. SYBR gold-stained samples were detected in the FITC channel using filters with excitation 455 - 485 nm/emission 510 - 545 nm with an exposure time of 100 ms. DFHO-stained samples were detected in both the FITC channel with an exposure time of 200 ms and the Cy3 channel with an exposure time of 100 ms (excitation 512 - 552 nm/emission 565 - 615 nm). Peptide recruiting condensates, which contain 6-FAM labeled peptides, were imaged using an inverted microscope (Nikon Ti2-E) using a blue LED source (488 nm), and a FITC/GFP filter cube (em 520 nm); exposure time was set to 500ms, except the for the samples including 3sv2-TAT nanostars and TAT peptide, in which exposure was set at 100 ms. Slides were prepared by adhering a square of parafilm with a 3 cm diameter circle cut out in the center to a coverslip (48393-070, VWR). This parafilm provided a small z distance to alleviate the disruption of the condensates. 2.5μL of the sample was placed in the cut-out and covered by a second coverslip (48366-227, VWR).

**Image enhancement** All black-and-white fluorescence images in figures were processed in FIJI (ImageJ) using a custom macro implementing the following pipeline: calibration, contrast enhancement (10% pixels saturated), denoising via Gaussian Blur (sigma = 1.5), and background subtraction with rolling ball method (radius = 1, for extremely large droplets radius = 10). Images for peptide recruitment were thresholded with the same LUT window (11000, 18000) (16-bit) for the brightfield channel and the same LUT window for TAT-6FAM (2500, 20000) or P22 N-6FAM (15000, 60000). Brightfield channel images were inverted to a black background before merging.

**Size analysis** Size analysis of each condition was done using data from 3 experimental replicates unless otherwise specified. Each experiment with purified RNA includes 7 fields of view (FOV) (a total of 21 images per analysis); each cotranscription experiment includes FOV = 11 (a total of 33 images per analysis); each cell-free cotranscription experiment includes FOV = 11. Images affected by strong background noise, defocusing, or overlapping FOV were discarded. To avoid image bias selection, we included the first 7 processable images collected in each experiment. All images were first enhanced, thresholded, and processed with a binary mask using a FIJI macro similar to the one described above. We set contrast enhancement with 0% saturation, Gaussian Blur with sigma = 2, and rolling ball radius = 0.5 to prevent the selection of artifacts due to defocusing or background noise. Conversion to mask was done via auto thresholding (Moments method). For condensate number analysis, objects smaller than 6 $px^2$ were considered noise and excluded. Droplet numbers were autogenerated by FIJI and recorded. Masks were saved as TIFF files for droplet size analysis.
Droplet size information was extracted as Chord Length Distribution (CLD) from the binary masks using a Python3 script based on PoreSpy, particularly Porespy.filters.apply_chords, which relies on Scipy and Skimage. The selected options allow for adjacent chords (spacing = 1) and include objects touching the edges of the FOV (trim_edges = False).
For each binary mask, the script calculates chords along the x and y directions via the chord_counts function from PoreSpy. The resulting Numpy arrays are saved in NPY format via Numpy. Each sample's resulting chord lengths, along x and y and across all FOVs, are pooled together, converted to μm via the pixel scaling within ND2 metadata, and saved into new arrays. The mean ($\mu_{CLD}$) and standard deviation of resulting CLDs over time are calculated for each sample by combining all FOVs. The reported mean are calculated as mean(mean_sample1, mean_sample2, mean_sample3); the standard error was calculated as std(mean_sample1, mean_sample2, mean_sample3).

##### 1.4.3 Image analysis using coordinate angle mapping

To determine whether our nanostars including distinct fluorogenic aptamers produce condensates that mix or do not mix, we built pixel intensity histograms from fluorescence microscopy images (manuscript Fig. 4). This was done because upon DFHO staining, both Corn and Red Broccoli aptamers can be detected in the FITC channel and the Cy3 channel, albeit with varying intensity, since Corn aptamer $Em_{max}$ = 545 nm, and Red Broccoli $Em_{max}$ = 582 nm, [2]. Images were processed to obtain the intensity of each pixel within segmented regions of interest in each channel (FITC and Cy3) and computing the angle of their coordinates in a plane with an x-axis corresponding to the CY3 channel intensity and with a y-axis corresponding to the FITC channel intensity, as sketched in Fig. S14. A line was drawn from the projected point to the origin, and the angle between this line and the x-axis was referred to as the coordinate angle. Finally, we generated a histogram based on the coordinate angles calculated from every pixel within the regions of interest.

It is important to note that although both Corn and Red Broccoli rely on the formation of G-quadruplexes to bind their fluorophore, Red Broccoli forms an intramolecular core-binding region, while Corn forms homodimer where fluorophores bind dimer interfaces[2,9]. This aptamer-directed dimerization is present in addition to designed kissing-loop-directed interactions, and results in the formation of structures other than spherical droplets for Corn-appended nanostars (**Fig. S15**, indicated by white arrow). These structures can be distinguished based on their non-spherical shape and were excluded from histogram generation according to their region labels.

As a control, we prepared separate samples including condensates from nanostars including either Corn or Red Broccoli, as shown in the upper panel in **Fig. S15**. The expectation is that if the nanostars are orthogonal, the histogram peaks measured when Corn and Red Broccoli condensates are produced in the same sample would be aligned with those of the separately annealed controls. If any mixing occurs due to the nanostar interactions, then a shift of the peaks (indicating mixing with a specific stoichiometry) or a uniform distribution (indicating mixing following a random stoichiometry) would be observed.

These experiments were done in triplicate, each with FOV = 10, for a total of 30 images. We used a bash script to randomly select 10 FOV in the set of images we collected and then execute image segmentation and pixel intensity extraction. Subsequent processing was performed using a Python3 script. For each FOV, we generated a mask by applying a Gaussian blur (sigma = 5) and gamma enhancement (gamma = 0.5) to the CY3 channel image, thresholding it using the Otsu method, skeletonizing it to generate seeds, and growing it using a watershed transformation. For each pixel within the masks, pixel intensities for CY3, FITC channels, and the region labels were extracted into three columns and saved in a .csv file. We manually excluded regions with unfocused droplets and non-spherical, cloudy condensates. Finally, we used a Python script to read and concatenate the data of 10 FOV into a single .csv file. Calculating coordinate angles and histogram plotting is described below (Data visualization).

##### 1.4.4 Data visualization

Violin plots were generated using a Python3 script based on Seaborn. For three NPY files storing chord length information of three repeats, the script loads the files, calibrates chord length into µm, and pools them into one array as the input dataset for violin plots. Cut = 0 was applied to ensure that the kernel density estimation doesn’t exceed the data range. As for kernel density estimation for the violin plots, data was normalized to have the same area of the violins (scale = ‘area’).  The coordinate angle histogram was generated using a Matlab script. The script loaded .csv files in which we recorded the pixel intensity of FITC and Cy3 channels of regions of interest (condensates). The background was eliminated by subtracting the smallest value of each channel. Each row of the resulting array contained the background-subtracted pixel intensities of both channels for each pixel within the droplet region. The coordinate angle for each pixel was calculated as arctan (FITC intensity/Cy3 intensity). Histograms were normalized to have the unitary area.

### 2 Supplementary Experiments and Text

#### 2.1 Multi-stranded nanostars (Caltech)

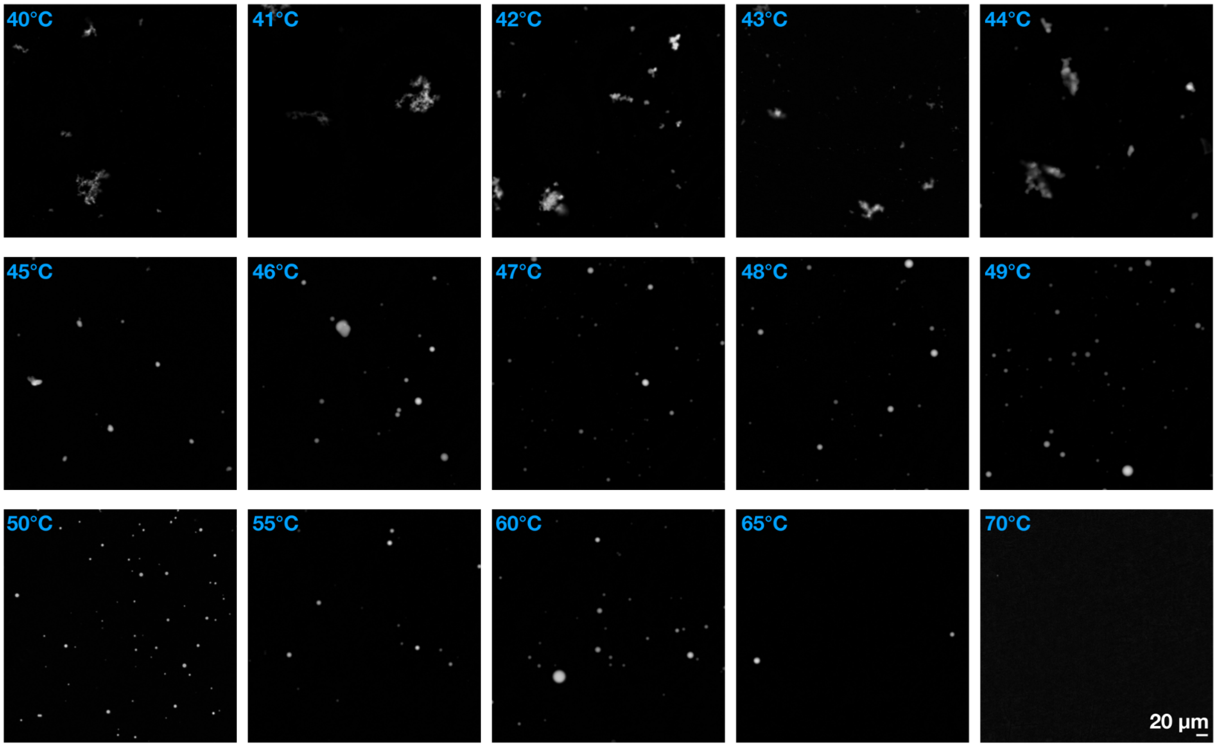


**Figure S1. 12 hour incubation over variable temperatures of multi-stranded condensates.** Multi-stranded condensates (4m5) were annealed with our annealing protocol that includes a melt phase (70°C for 10 min) and hold phase (temperature between 40°C-70°C 12h) protocol. Strands were annealed in our assembly buffer (40 mM HEPES/100KCl/500 mM NaCl). Samples were stained with SYBR Gold and imaged.

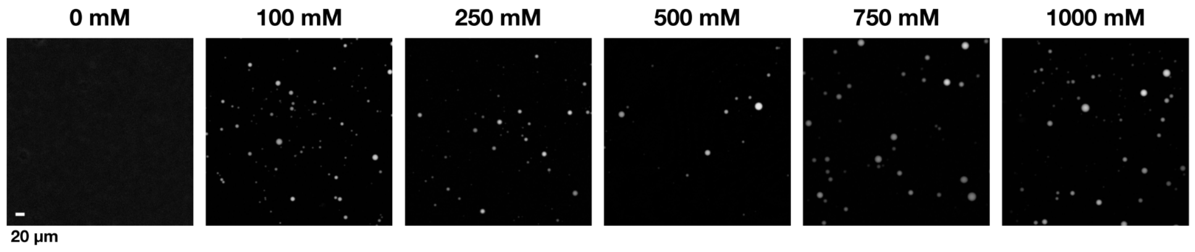


**Figure S2. Potassium titrations of multi-stranded condensates.** Nanostars (4m6) were assembled in 40 mM HEPES and a variable KCl concentration (Figure S3). All samples underwent a thermal treatment of a 70°C denaturing steep for 10 minutes, then incubated at 50°C for 12 hours, then a quick cool to room temperature. Liquid-like condensate formation begins at 100 mM KCl and maintains droplet-like morphology through 1000 mM KCl. Samples were stained with SYBR Gold and imaged.

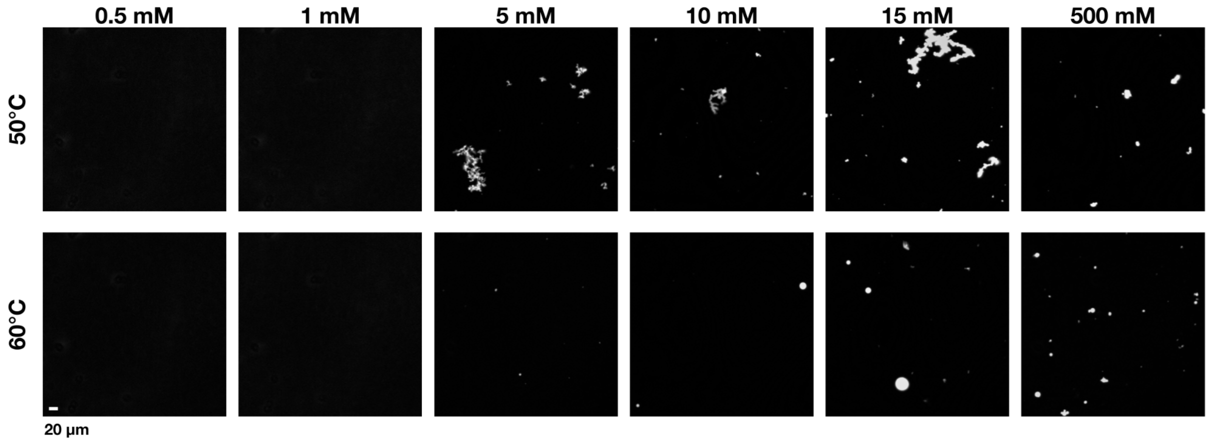


**Figure S3. Magnesium titrations of multi-stranded condensates.** Nanostars (4m6) were assembled in 40 mM HEPES and a variable Magnesium concentration. Under the protocol using equimolar purified RNA strands that form an RNA motif via a thermal treatment of a 70°C denaturing step for 10 minutes, then incubating at 50°C for 12 hours, then a quick cool to room temperature, the addition of Magnesium causes gel formation, starting at 5 mM. As expected, the addition of Magnesium changes the melting temperature of the condensates, in which we observe droplet-like assemblies when we raise to a 12 hour incubation step from 50°C to 60°C around 15 mM Magnesium. Samples were stained with SYBR Gold for imaging.

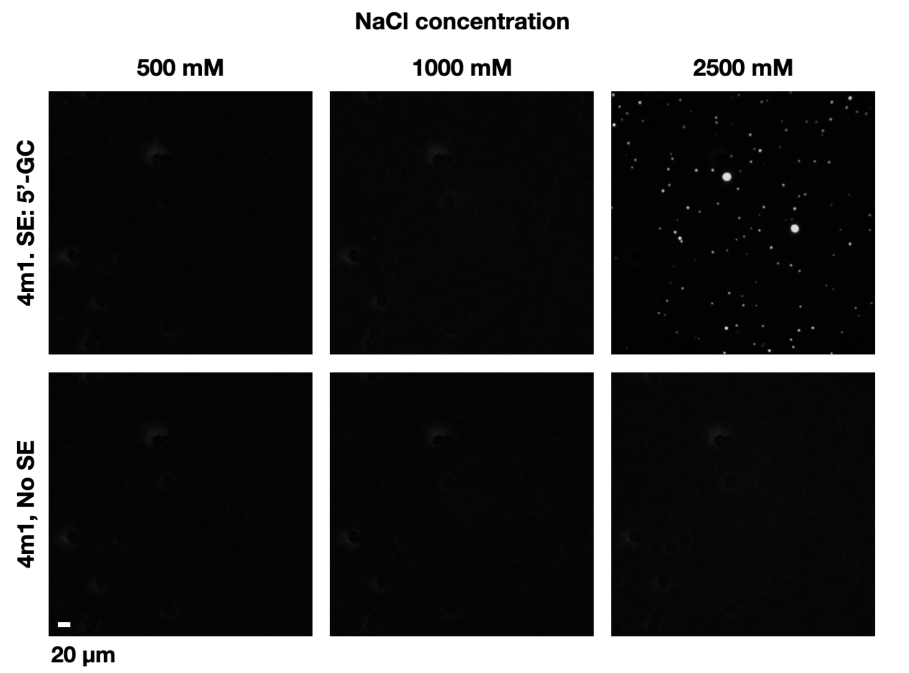


**Figure S4. NaCl titrations using multi-stranded nanostars with and without 2nt sticky ends.** Nanostar with 2nt 5'-GC (4m1) required 40 mM HEPES/100 mM KCl and increased NaCl concentration to form via a thermal treatment of a 70°C denaturing step for 10 minutes, then incubating at 50°C for 12 hours (Figure S5). RNA nanostars without sticky-ends (4m1) do not form in comparable buffer and temperature conditions. Samples were stained with SYBR Gold for imaging.

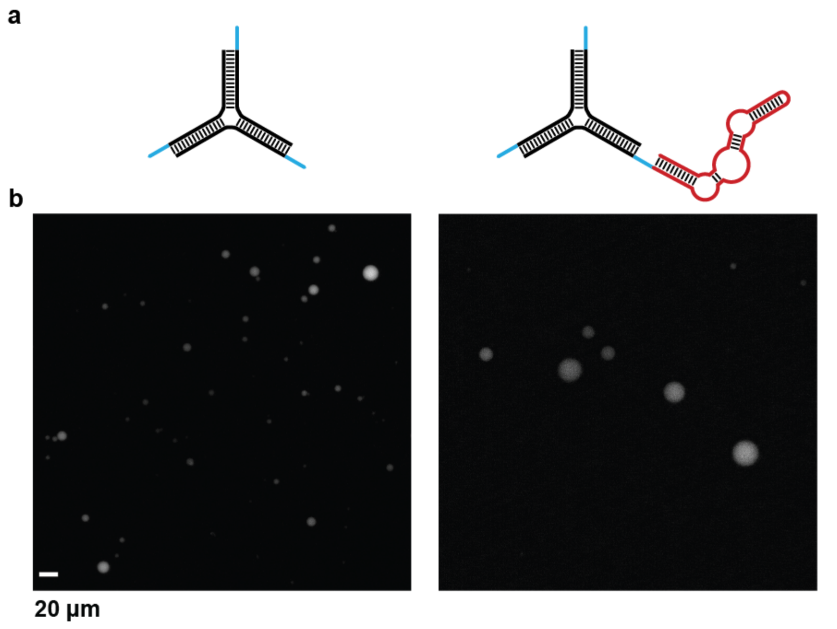


**Figure S5. Three-arm multi-stranded motifs form condensates.** Left, nanostar 3m1 modified from 4m6 (GCUAC), assembled in 40 mM HEPES, 100mM KCl, and 2.5M NaCl. Right, the same motif including the Red Broccoli aptamer, Cy3.
All samples underwent a thermal treatment of a 70°C denaturing steep for 10 minutes, then incubated at 50°C for 24 hours, then a quick cool to room temperature. Left, unmodified condensates were stained with SYBR Gold and imaged using FITC channel.

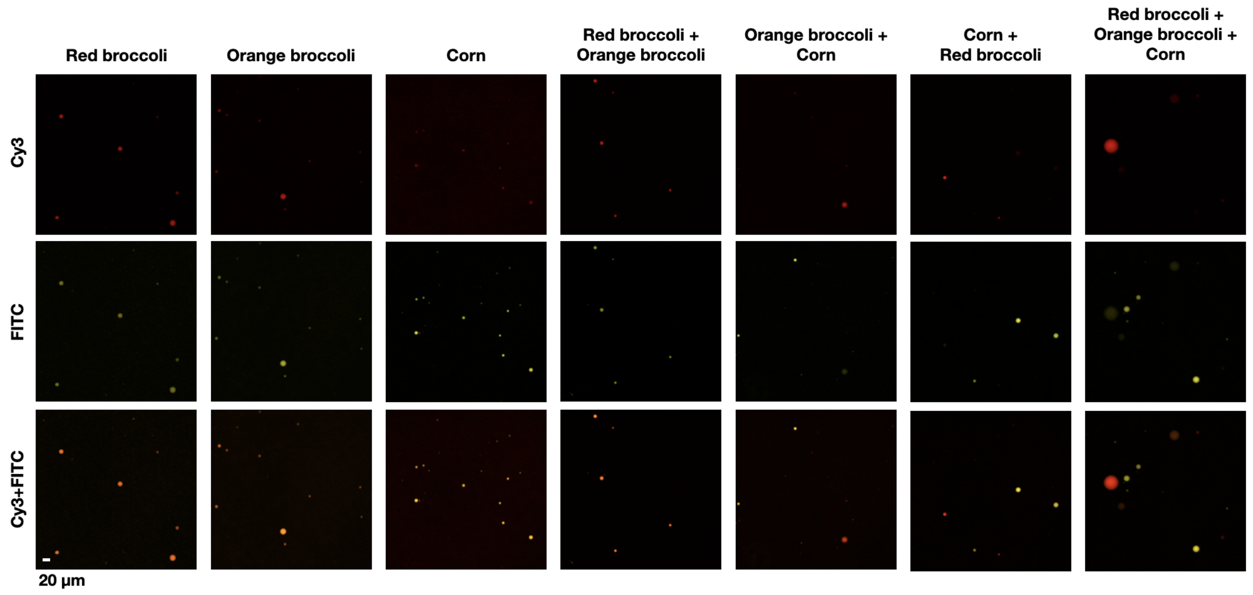


**Figure S6. Condensates with fluorogenic aptamers.** RNA nanostars 4m3, 4m5, and 4m6 with Orange Broccoli, Corn, and Red Broccoli aptamers respectively, formed as single, double, and triple condensate systems in 40 mM HEPES, 100 mM KCl, 500 mM NaCl. Condestes were thermally treated by a quick 70°C melt followed by a 12h-long hold 50°C. Condensates were imaged using single filter channels Cy3 and FITC and showing the resulting overlay of filters Cy3 and FITC.

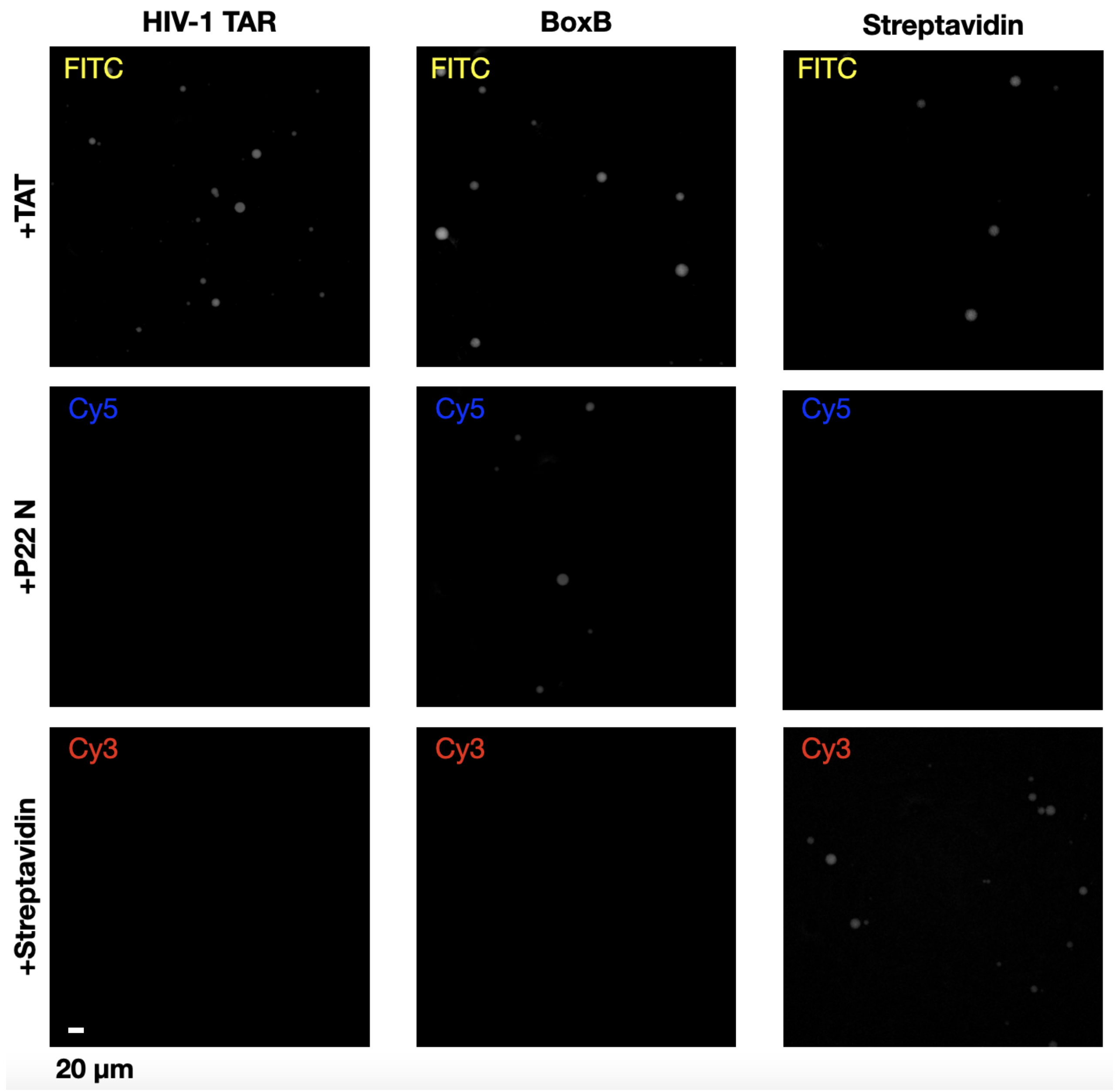


**Figure S7. Recruitment of peptides and protein of multistranded condensates.** Fluorescence microscopy data of multistranded condensates 4m6 with TAR RNA, 4m5 with boxB RNA, and 4m3 with streptavidin binding aptamer in the presence of individual targets TAT peptide labeled with 6FAM, P22 N peptide labeled with AF647, and streptavidin protein labeled with AF555. The positive charge of TAT peptide due to the rich arginine content causes indiscriminate binding to negatively charged RNA. Scale bar, 20 μm.

#### 2.2 Three-arm single-stranded nanostars (UCLA)

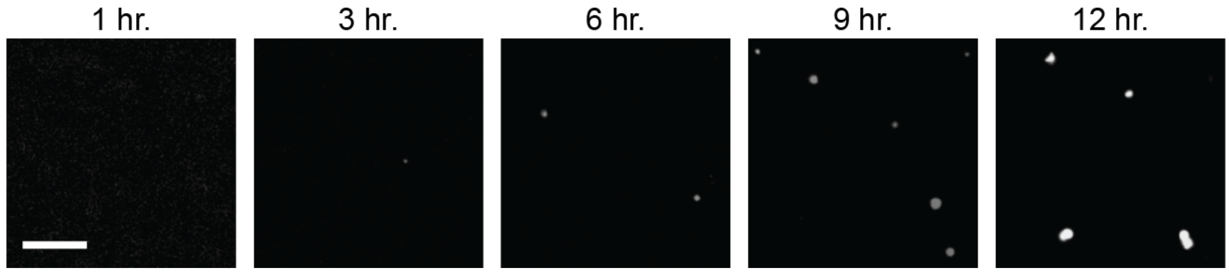


**Figure S8. Condensate growth during incubation at 50°C.** We tracked growth of our three arm single stranded nanostar 3sWT-stem1 during the hold step. The annealing mixture was aliquot into five samples, initiated simultaneously, and imaged at different time points. Condensates were stained with SYBR Gold and imaged. Images are enhanced for brightness and contrast with the same treatment for displaying. Scale bar, 10 µm.

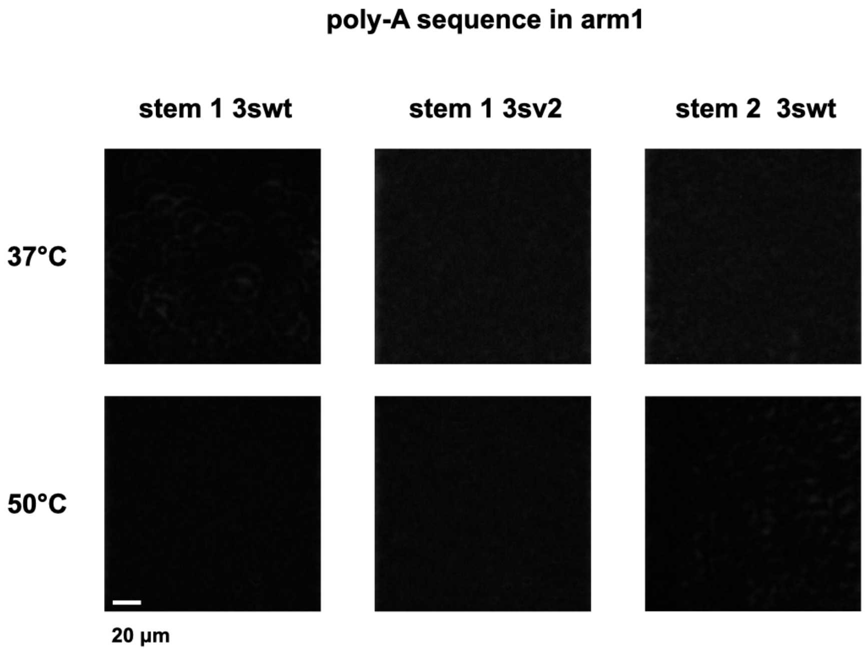


**Figure S9. Single-stranded nanostar variants with one of the three WT KLs replaced with the poly-A sequence formed no condensate.** After the melt step, nanostars were incubated at 37°C or 50°C for 12h and cooled to 20°C before imaging. Samples were stained with SYBR Gold for imaging. Scale bar, 20 µm.

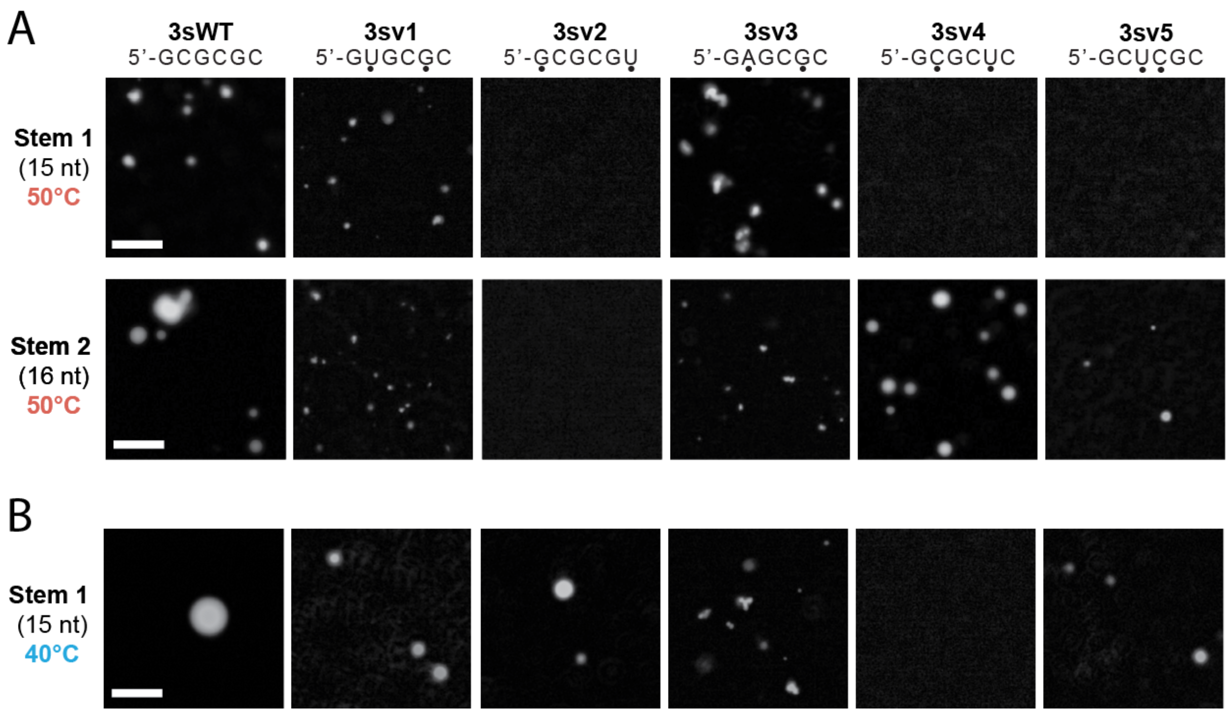


**Figure S10. Overview of fluorescence microscopy images for single nanostar designs. A)** Condensates assembly from two types of stems and six types of self-complementary kissing-loops that had no (3sWT) or two (all variants) mismatches (indicated by dots) with a 50°C hold. Stem 1 and 2 have different lengths and sequences. **B)** Condensates assemble from Stem 1 variants with a 40°C hold. Dropping of holding temperature facilitates condensation evidenced by appearing of condensates for 3sv3 and 3sv5, and increasing of sizes for 3sWT and 3sv1. All variants were assembled from purified RNA using our assembly buffer and the melt and hold annealing protocol (10 min at 70°C followed by 12 h hold at specified temperature), and stained with SYBR Gold for imaging. Scale bar, 10 μm.

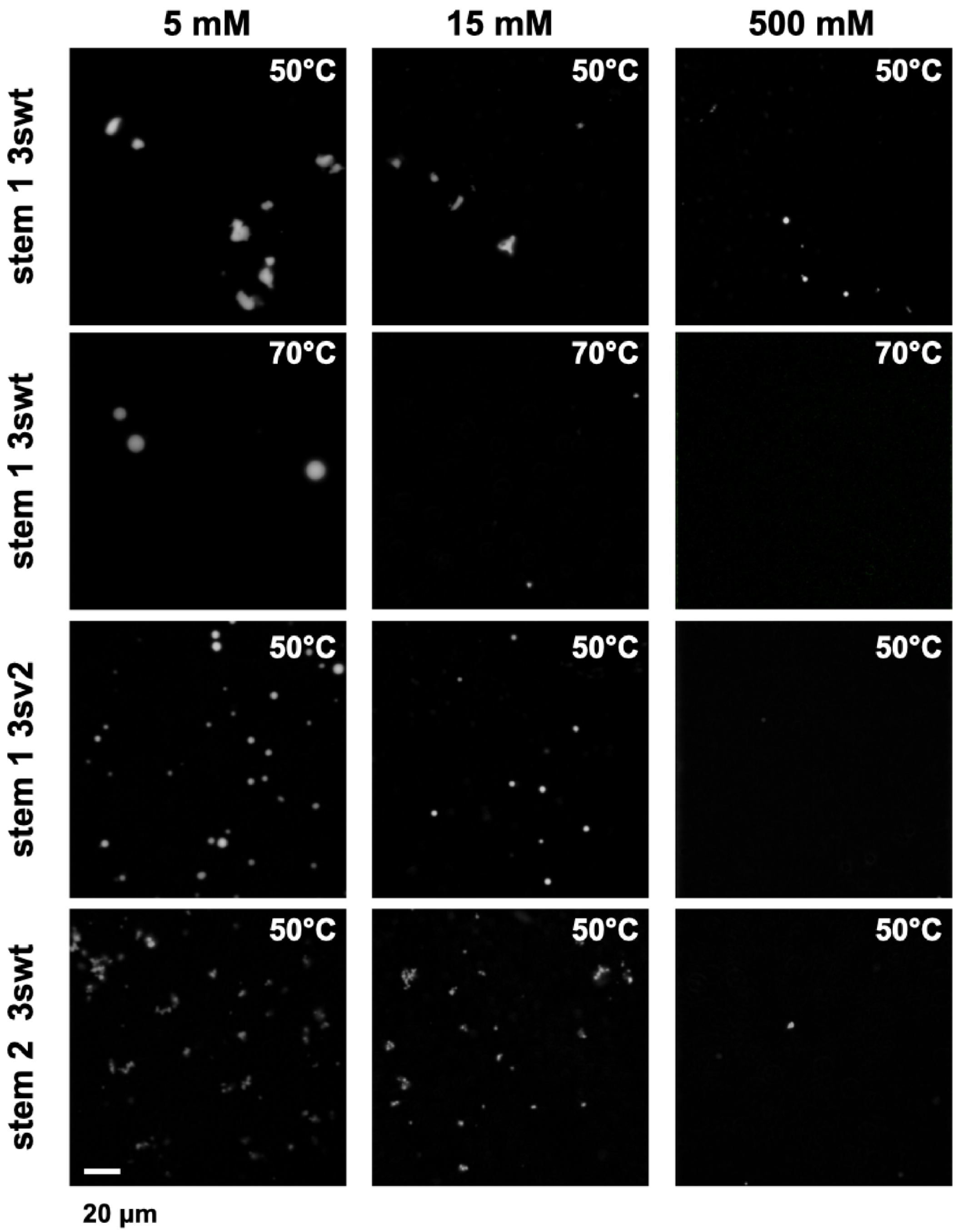


**Figure S11. Condensate formation at different magnesium conditions.** Fluorescence micrographs showing condensate morphology of three motifs in the presence of 40 mM HEPES and a variable MgCl2 concentration. Condensates were annealed at 50°C for 12h and cooled to 20°C before imaging. Samples were stained with SYBR Gold for imaging. Scale bar, 20 µm.

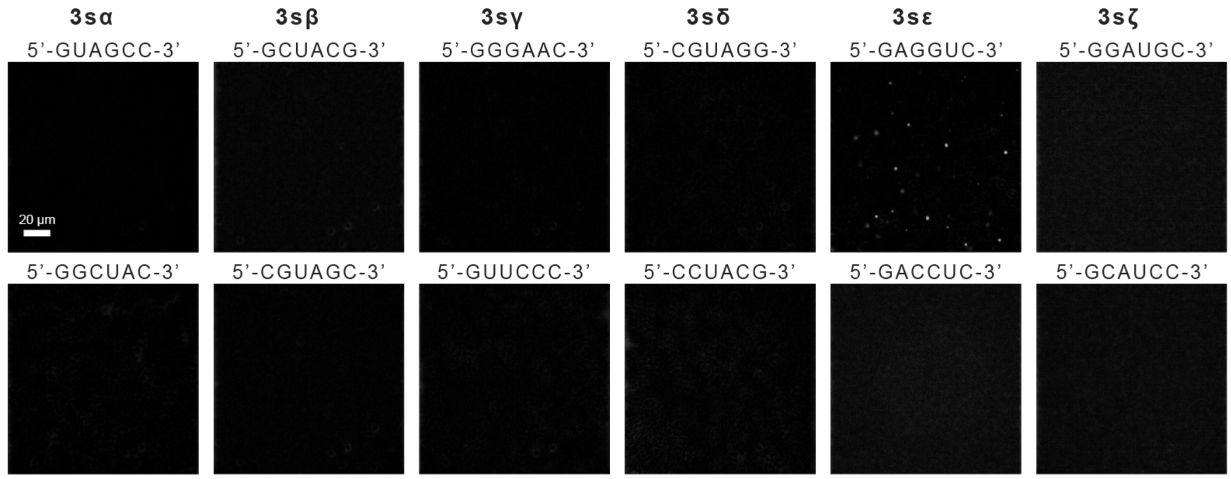


**Figure S12. Condensates do not form when annealing individual components of the two nanostar designs.** Motifs have identical stems, and these results indicate that stem-stem interactions do not determine condensate formation. Only variant 3sε1 form condensates; the KL of this variant includes two guanine pairs surrounded by a total of four complementary base-pairs, confirming that mismatches within kissing loops can be tolerated and still yield condensates. However, condensation is sensitive to the type of substitution and to the position of mismatches within kissing loops (see Fig. S10). Scale bar, 20 µm.

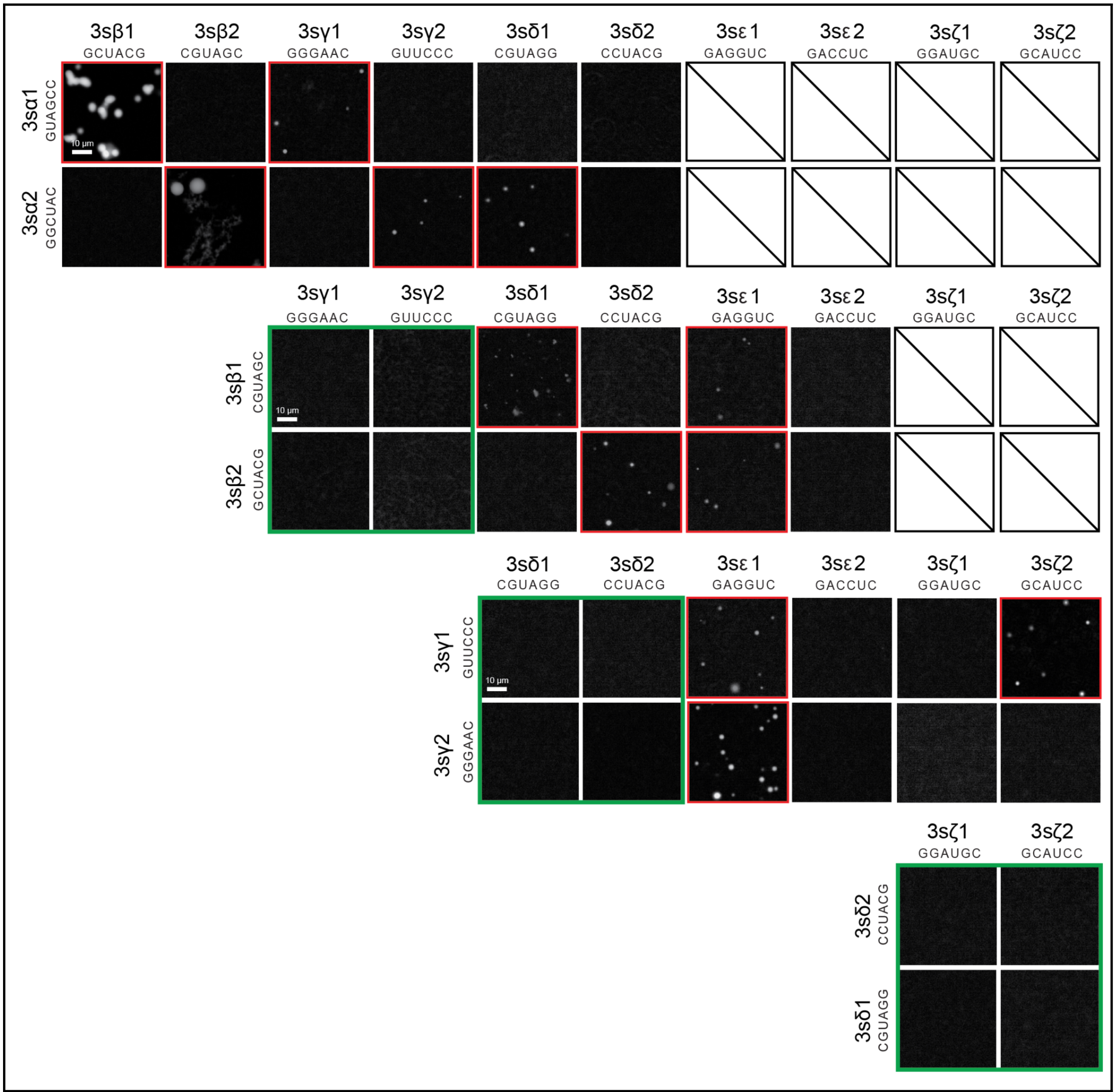


**Figure S13. Cross orthogonality between two nanostar condensates.** Each box includes representative microscopy images of samples prepared by annealing (melt and hold at 50°C) nanostar motifs that are part of two nanostar generating condensates. The motif name and sequence are marked to the left of each row and at the top of each column. Pairs that produce condensates are highlighted in red. Pairs 3sβ/3sγ and 3sγ/3sδ are fully orthogonal (in green). Pair 3sβ and 3sγ are used for orthogonality demonstrations in the main paper (Fig. 4C). Pair 3sζ fails as it generates gel-like networks which do not match our goal of generating liquid-like droplets (see Fig. 3G in the manuscript). Pair 3sε fails as nanostar 3sε1 yields condensates on its own (Fig. S12). Scale bar, 10 µm.

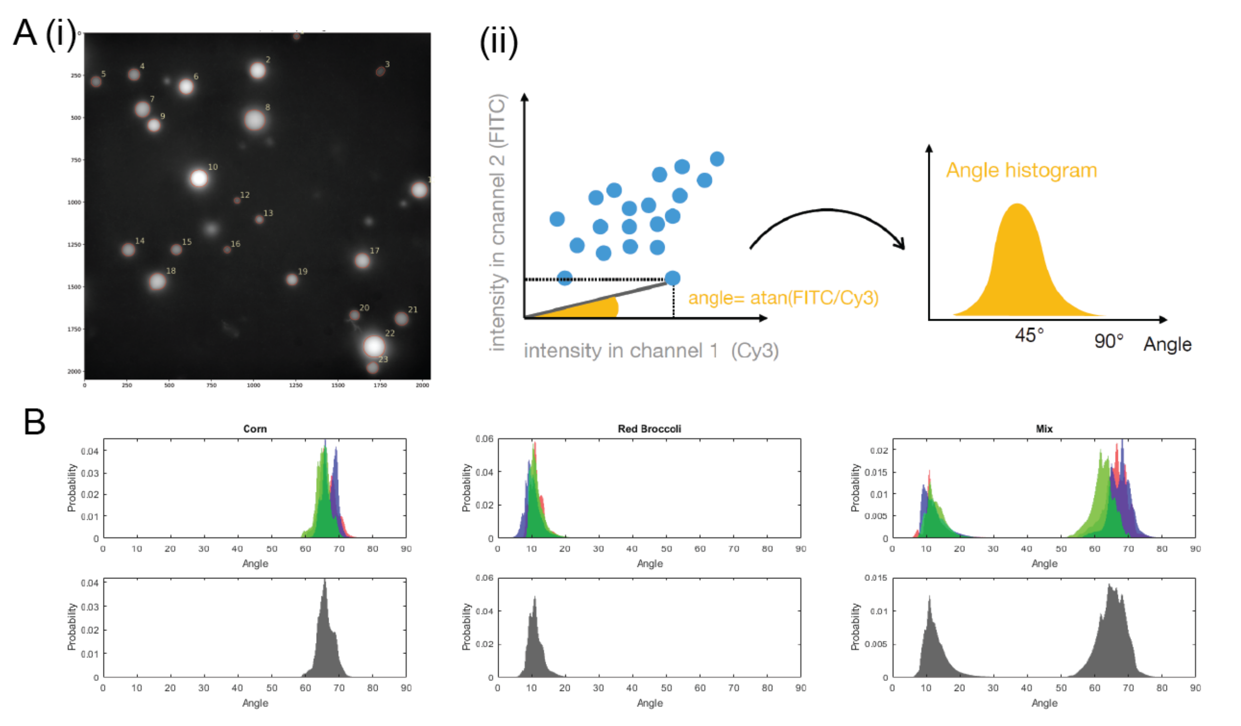


**Figure S14. Coordinate angle mapping workflow and results for each sample.** (**A**) (i) Example of an overlay of fluorescent image and labeled regions that were drawn from corresponding binary mask. We extracted pixel intensities in the CY3 and FITC channels within these labeled regions (circled in red). (ii) For each pixel, a coordinate angle was calculated as arctan (FITC_intensity/CY3_intensity). A histogram of coordinate angles was generated for each sample (FOV = 10). (**B**) Histograms of N = 3 samples (top panel, in red, green, and blue) and sum of the three (bottom panel, in gray) for 3sγ_Corn, 3sβ_Red Broccoli, and one-pot annealing of the two. The skewing towards smaller angles (for Corn) and bigger angles (for Red Broccoli) were due to saturated pixels in the FITC channel (for Corn) or CY3 channel (for Red Broccoli). Total RNA concentration doubled in mixed annealing, resulting in bigger droplets and more saturation, thus increasing the skewing of peaks towards the middle.

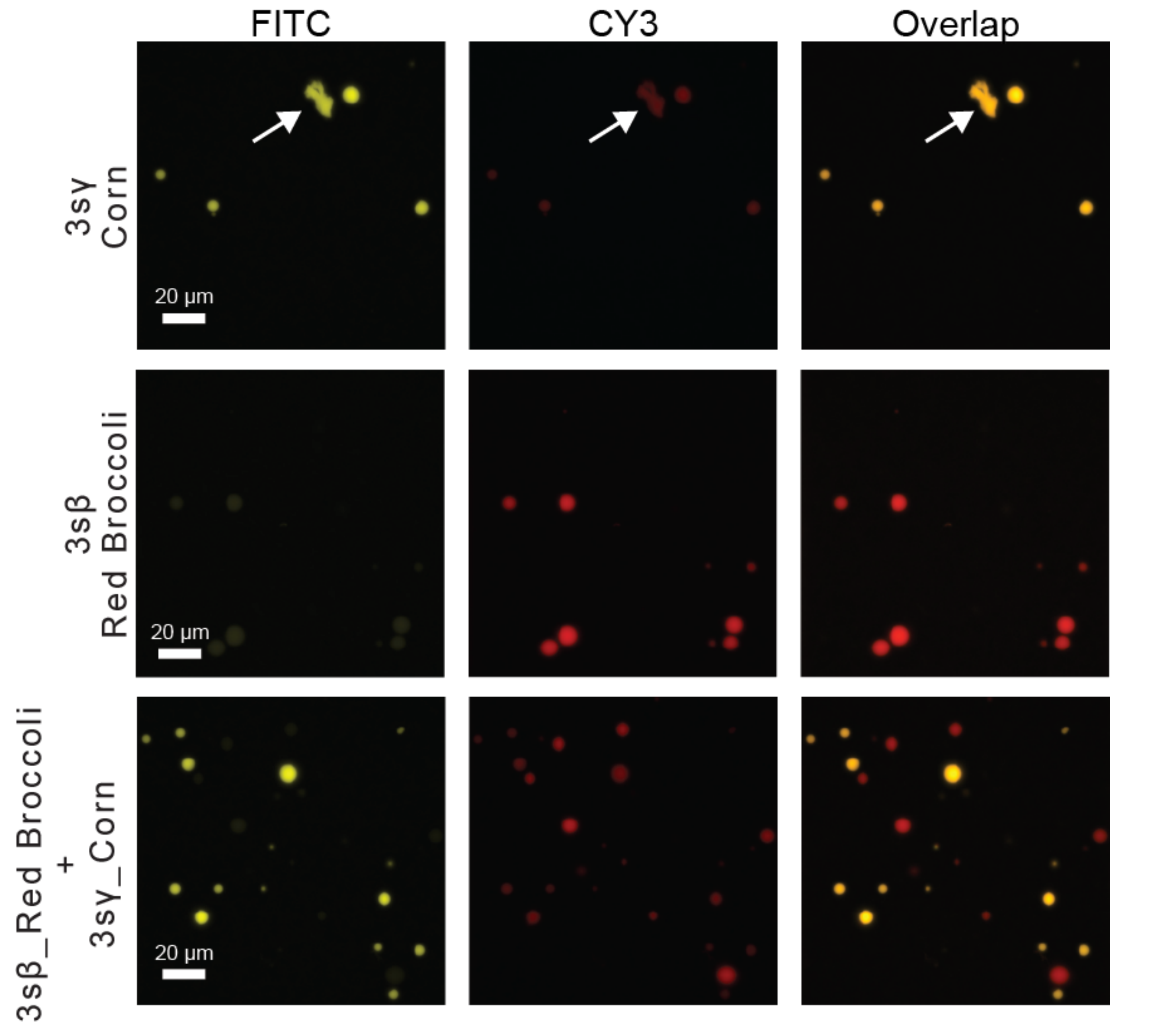


**Figure S15. Example images of individually and simultaneously annealed two nanostar condensates that include fluorogenic aptamers.** Rows 1 and 2 from the top are individually annealed nanostars, row 3 shows example images of simultaneously annealed nanostars. The nanostars include 25% of aptamer-appended strands. White arrows point at non-spherical condensates whose formation may be driven by dimerization of Corn aptamers.

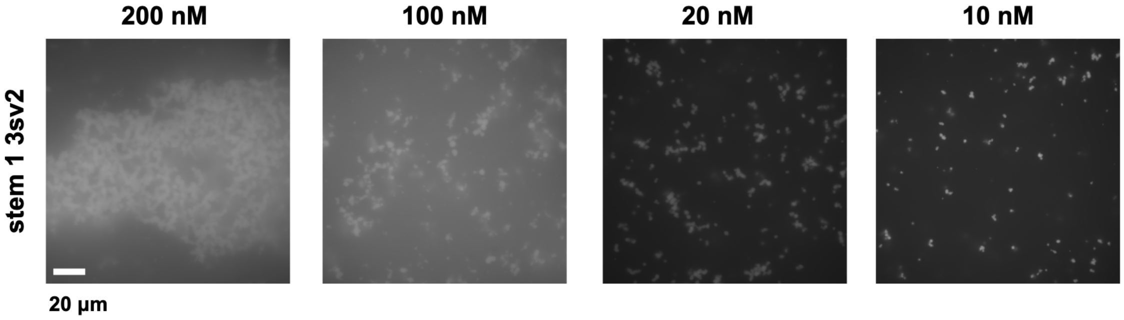


**Figure S16**. **DNA template titration for co-transcription experiments.** Fluorescence micrographs at 2h timepoint showing cotranscriptional condensate formation with a variable final DNA template concentration. Scale bar, 20 μm.

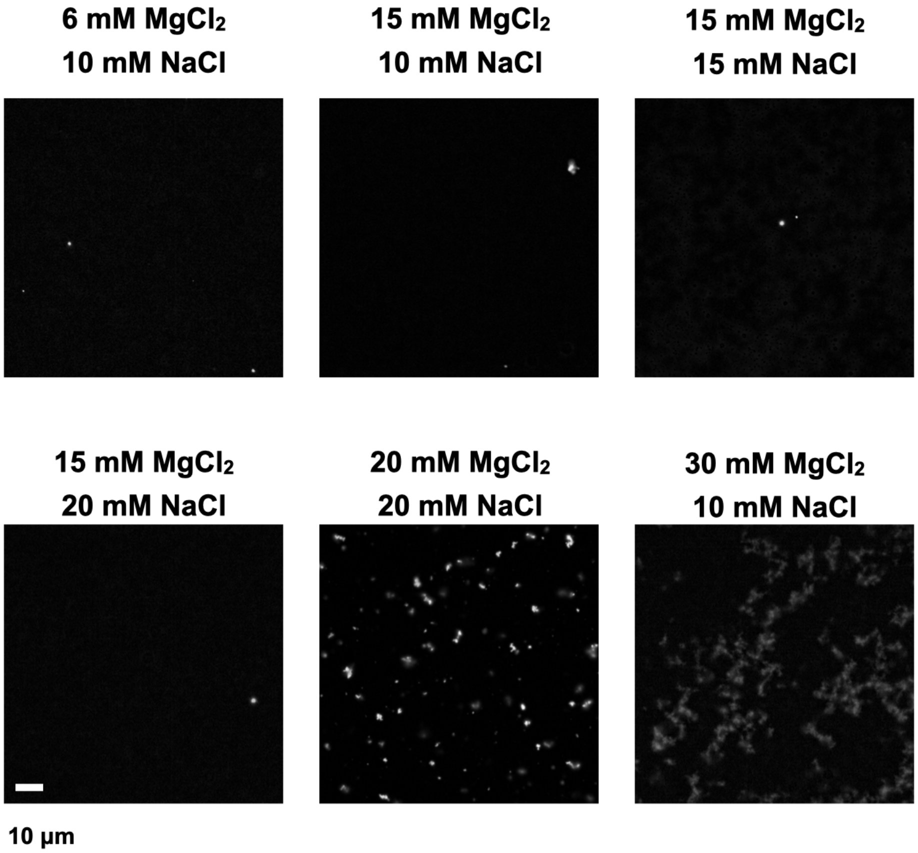


**Figure S17. Magnesium and sodium titration for co-transcription experiments of the two-nanostar system (**3sβ**).** Fluorescence micrographs at 1h timepoint showing cotranscriptional condensate formation with various MgCl2 and NaCl concentrations. Each template is added at 10 nM. Reaction conditions are consistent with those listed in section 1.3.2. Scale bar, 10 μm.
